# Characterization of Single Ribosomes and Virus Particles with Large-Diameter Perfringolysin O Nanopores

**DOI:** 10.64898/2026.09.16.752082

**Authors:** Andela Vracar, Veneta Salyahetdinova, Sandor Balog, Alessandro Ianiro, Anasua Mukhopadhyay, Michael Mayer

## Abstract

Existing biological nanopores are often too narrow to accommodate large proteins and biomolecular complexes in their native folded states, precluding the analysis of megadalton assemblies. Here, we report the self-assembly of the cholesterol-dependent cytolysin Perfringolysin O (PFO) into stable transmembrane nanopores composed of 46 ± 9 monomers, with an inner pore diameter of 28.5 ± 5.6 nm and a length of 9.8 nm. We demonstrate that despite the large size of the pore assembly, PFO pores exhibit a stable open-pore current with a high signal-to-noise ratio, making them suitable for resistive-pulse recordings. Moreover, PFO nanopores enable accurate, calibration-free, and label-free sizing of individual proteins, multi-protein complexes, and viral particles across a large molecular-weight range spanning 50 kDa to 3.4 MDa. The exceptionally large diameter of these pores enables, for the first time, resistive-pulse-based characterization of intact virus particles and ribosomes with a biological nanopore. Specifically, we determined the volume, shape, and diameter of complete capsids of recombinant adeno-associated virus serotype 2 (rAAV2) as well as capsid fragments. Finally, simultaneous analysis of molecular volume and shape resolved intact 70S ribosomes from their dissociated 30S and 50S subunits in a mixture. By extending biological nanopore sensing to single-particle characterization of large protein complexes well beyond the reach of existing pores, this approach introduces PFO nanopores as a versatile platform for label-free, single-particle analysis in solution.

## Introduction

Ionic current recordings through a nanoscale pore in an insulating membrane provide a label-free method of detecting and characterizing individual macromolecules at the single-molecule level.^1–4^ Characterization of individual analytes is achieved by inserting a nanometer-scale pore into an insulating membrane that separates two chambers filled with electrolyte solution. When a voltage is applied, diffusing analytes are driven through the electrolyte-filled pore by electrophoretic force, electroosmotic flow, or both. The presence of the analyte in the pore lumen produces a measurable reduction in ionic current, known as a resistive pulse. The amplitude of the current blockade and the modulation of the intra-event current during this pulse directly reflect the size, shape, and rotational diffusion coefficient of the analyte molecules in a label-free fashion.^5–7^

Nanopore sensing platforms are broadly classified into solid-state^8^, biological^9^, and hybrid systems.^10^ Solid-state nanopores fabricated in silicon nitride, graphene, and aluminum oxide membranes offer the ability to prepare diameters large enough to accommodate megadalton-scale biomolecular assemblies, including intact viruses, viral capsids, ribosomes, and large protein complexes.^11–18^ Despite these advantages, solid-state nanopores present several limitations for the analysis of complex biomolecular assemblies. Solid-state nanopores are prone to long-term instability, exhibiting changes in pore diameter and geometry due to slow etching and surface interactions with the recording electrolyte. To minimise the effect of slow etching and to prepare robust membranes, solid-state nanopores with diameters exceeding 20 nm are typically fabricated in relatively thick (>20 nm), free-standing membranes. This approach increases the length of nanopores, thereby reducing resistive pulse amplitudes for the same analyte particles compared with short nanopores. Moreover, free protein translocation through solid-state nanopores typically yields many resistive pulses with short dwell times, often below the measurement’s time resolution, or provides limited sampling of particle orientation per resistive pulse. The opposite extreme of long dwell times, resulting from non-specific protein adsorption, is also problematic, as it can lead to biased sampling of particle orientation or pore clogging.^19–22^

Biological nanopores in microscale planar lipid membranes^23^ in comparison, typically exhibit lower electrical noise ^24^, reduced non-specific adsorption, and a stable, well-defined geometry under appropriate experimental conditions.^25^Their pore size is, however, constrained by the molecular structure of the protein pore or by the self-assembly properties of the pore-forming proteins. At present, the largest biological nanopores are comprised of complement component 9 (poly(C9) pores),^26^ which forms cylindrical pores approximately 10 nm in diameter and 13 nm in length, and Pneumolysin (PLY), a cholesterol-dependent cytolysin that assembles into pores approximately 20 nm in diameter and 9.5 nm in length.^27^ The inner diameters of these pores substantially exceed the dimensions of previously reported nanopores, such as the two-component pleurotolysin pore, PlyAB, with a *cis* pore opening of 10.5 nm, a *trans* pore opening of 7.2 nm, and an internal constriction of 5.5 nm.^28^ So far, no biological nanopores with diameters larger than approximately 20 nm have been reported.

To overcome this size limit, the work presented here introduces a biological nanopore self-assembled from Perfringolysin O (PFO), a cholesterol-dependent bacterial cytolysin (CDC)^29^ that belongs to the family of pore-forming toxins (PFTs), secreted by *C. perfringens* as a water-soluble monomer.^30^ PFO binds to cholesterol-containing membranes and spontaneously self-assembles into large pores through a multistep mechanism involving membrane binding of the monomer, oligomerisation to form a ring-shaped pre-pore complex and insertion of the pre-pore’s β-hairpins into the lipid membrane to form the pore.^31–33^ Previous studies have shown that up to 50 PFO monomers self-assemble into cylindrical pores with inner diameters ranging from 25 to 30 nm.^33^ The combination of the large diameter and approximately cylindrical geometry^34,35^ makes PFO pores particularly well-suited for nanopore experiments with natively folded protein analytes because the electric field along cylindrical nanopores can be assumed constant. A constant electric field along the length of the nanopore is an important prerequisite for accurate determination of protein shape and volume.^7,21^

To demonstrate the applicability of PFO for characterizing large biological assemblies, we first revealed their exceptionally large and stable inner diameter of 28.5 ± 5.6 nm, which, to the best of our knowledge, is the largest lumen diameter reported for a biological nanopore used for resistive pulse analysis to date. We then employed these ultra-large pores to investigate the volume and shape of a range of folded proteins with molecular weights spanning from 50 to 480 kDa. Finally, we demonstrated the capability of PFO pores to characterise the size and shape distributions of recombinant AAV2 virus-like particles (rAAV2 VLPs), intact 70S ribosomes, and their constituent subunits on the single-particle level.

## Results and Discussion

### Self-assembly of PFO monomers to large nanopores and their characterization

To prepare PFO stock solutions for nanopore experiments, we combined previously reported protocols for PFO pore assembly on cholesterol-containing unilamellar liposomes with existing protocols for poly C9 nanopores and PLY nanopores.^26,27,37–39^ To this end, we purified PFO monomers (MW = 55 kDa) and pre-incubated them with DTT and amphipol A8-35. The structure of the PLY pore (PDB: 2BK1) in **Figure 1A** was obtained by fitting the α-carbon trace of perfringolysin O into a cryo-EM density map, as no high-resolution structure of Perfringolysin O in its pore-forming conformation is currently available; both PLY and PFO share conserved structural architecture within the cholesterol-dependent cytolysin family.^36^ To assemble and insert PFO nanopores in planar lipid bilayers, we added the pre-incubated solution at a final PFO concentration of 8.1 µg/mL to the *cis* compartment of the recording chip (**Figure 1A**) with a planar lipid bilayer containing 30 mol % cholesterol and applied a potential of +100 mV to the *trans* side of the bilayer using two Ag/AgCl electrodes. We monitored pore formation *via* electrical recordings **(Figure 1A**). Typically, in a recording buffer at pH 7.5, PFO pores inserted into the bilayers within 12 min, causing a sudden, single-step increase in ionic current (**Figure 1B**). The distribution of the observed magnitudes of the current jump upon pore formation in **Figure 1C** reveals an inner diameter of the membrane-reconstituted nanopores of 28.5 ± 5.6 nm (mean ± SD, *N* = 55). These diameters correspond to 46 ± 9 PFO monomers (**Figure 1C**, **Supplementary Note 1**), in excellent agreement with cryo-EM results.^33^

**Figure 1.**
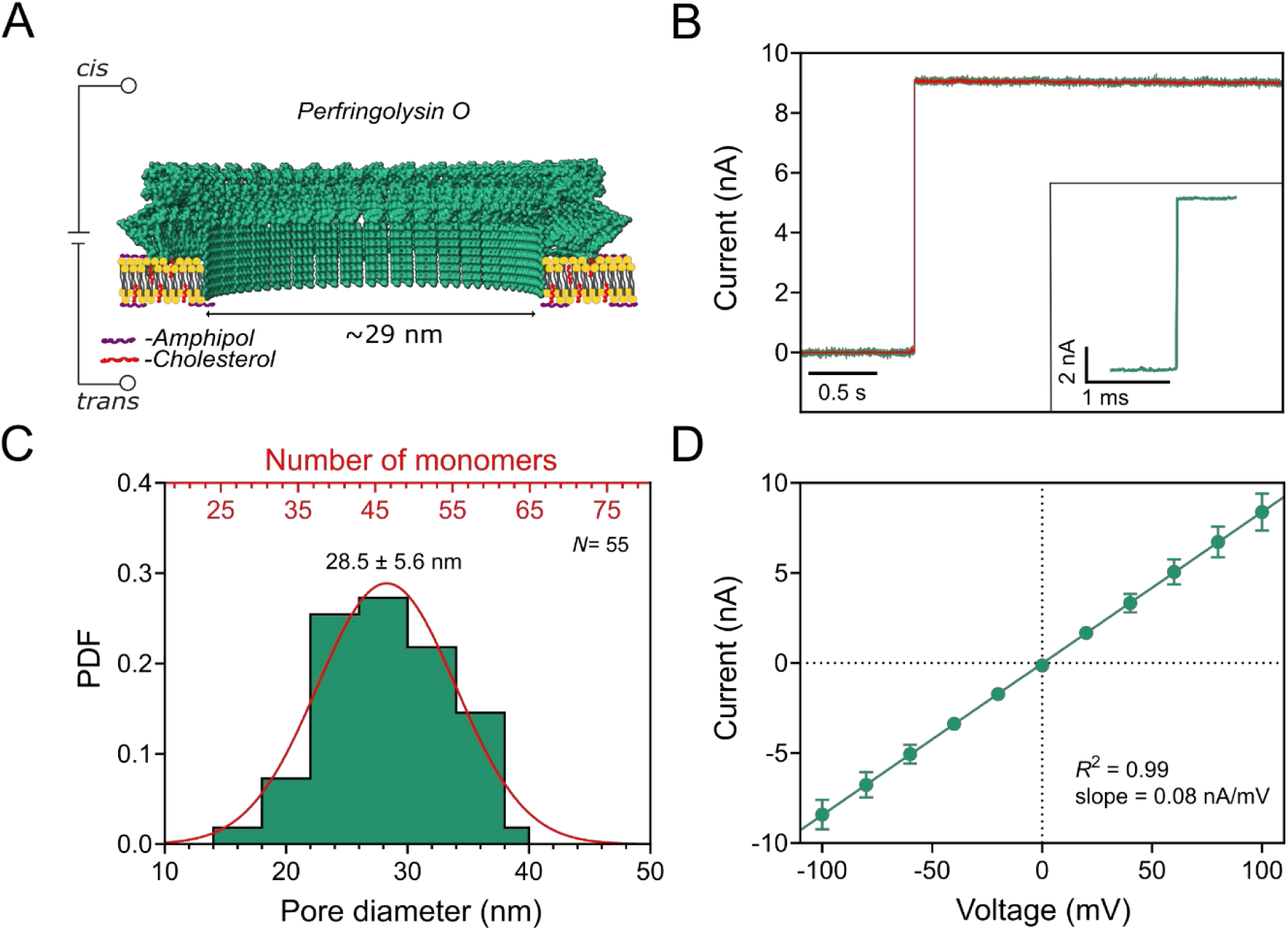
Characterization of self-assembled PFO nanopores in cholesterol-containing planar lipid bilayer membranes. (**A**) Schematic representation of a section of a PFO nanopore in a cholesterol-containing planar lipid bilayer. The representation of the PFO nanopore was obtained by fitting the alpha carbon trace of Perfringolysin O into a cryo-EM map of Pneumolysin (PDB 2BK1).^36^ (**B**) Current recording showing a single-step formation of a single PFO nanopore in a planar lipid bilayer under an applied potential of +100 mV. The data shown in green were sampled at the amplifier’s maximum rate (200 kHz) with no low-pass filter applied; the data shown in red were filtered with a Gaussian low-pass filter with a cutoff frequency of 10 kHz. The recording buffer contained 500 mM NaCl, 50 mM Tris-HCl (pH 7.5), and 0.2 μM amphipol. The inset shows the moment of pore formation with an expanded time axis and a 100 kHz Gaussian low-pass filter. (**C**) Distribution of the diameters of single PFO pores in cholesterol-containing planar lipid bilayers under an applied potential of +100 mV; pore diameters were calculated from their conductance, assuming a cylindrical pore shape and a pore length of 9.8 nm (see below). The top axis in (C) represents the estimated number of PFO monomers that corresponds to the determined values of pore diameters (see **Supplementary Note 1**). (**D**) Current-voltage (I–V) curve through a PFO pore under the same conditions as in (B). Error bars represent the standard deviations calculated from at least three repeats.

To investigate the effect of ionic strength on PFO pore formation, we conducted experiments using recording electrolytes with NaCl concentrations ranging from 0.05 to 1 M, buffered to pH 7.5 with 50 mM Tris-HCl, and containing 0.2 μM amphipol (see **Supplementary Figure S1**). Although we observed PFO formation across all tested ionic strengths, 500 mM NaCl yielded the most consistent formations and was therefore selected as the standard condition for all experiments. To assess the stability and associated noise characteristics of PFO nanopores, we compared the ionic current noise before and after pore formation in a recording buffer containing 500 mM NaCl, 50 mM Tris-HCl, pH 7.5, and 0.2 μM amphipol based on power spectral densities (PSDs). Upon pore formation, the PSD spectra showed only slightly increased noise in the low-frequency domain below ∼ 1 kHz (see **Supplementary Figure S2**) compared to the baseline current before formation, suggesting small-amplitude fluctuations of the pore volume of PFO pores in the time range of milliseconds. Over time, the electrical noise measured from a bilayer with an inserted PFO nanopore remained stable with excellent low-noise characteristics for resistive pulse recordings (**Figure 1B**). The rms current noise of the baseline current was 0.008 nA, and after pore formation, it was 0.026 nA at 10 kHz bandwidth.

For the characterisation of the shape of natively folded proteins with nanopores, the electric field inside the nanopore must be as uniform as possible^21^ therefore, the shape of the nanopore must be as close as possible to a cylinder. The approximated molecular structure of PFO pores in **Figure 1A** indicates that the pore’s shape is close to a cylinder, and AFM studies show that PFO monomers inside PFO pore assemblies are oriented perpendicular to the membrane surface. In addition, **Figure 1D** shows that the current-voltage relationship (I−V curve) of membrane-inserted PFO pores displayed ohmic behaviour in a range of ionic strengths from 0.05 to 1 M NaCl (see **Supplementary Figures S3,** and **S4** for inner pore diameter calculation at different salt concentrations). The absence of ionic current rectification (ICR) of PFO pores at relatively low ionic strength (50 mM) supports the AFM observation that the pore lumen does not have a conical shape, since conical nanopores with non-zero net charge in their lumen usually display ICR at low ionic strength. The absence of ICR is therefore consistent with a cylindrical pore lumen inside the lipid bilayer membrane, although it does not exclude the possibility of pores with cross-sections that may not be perfectly circular, such as ellipsoidal cross-sections. AFM studies of PFO pores in supported lipid membranes conducted by Czajkowsky *et al.* show that the cross-sections of most PFO pores approach perfect circles.^40^ The small fraction of PFO pores with ellipsoidal cross-sections, in most cases, had ratios between the major and minor axes smaller than 2:1. These moderate deviations from a perfectly circular cross-section of the pores suggest that, in the context of resistive pulse recordings, the approximation of a cylindrical pore shape remains justified, as these deviations do not yet exert a strong effect on the pore resistance and access resistance of the pore. The same AFM studies also show a significant fraction of arc-shaped PFO assemblies. ^40^ These arcs did, however, not appear to form pores through the planar lipid bilayer membranes we prepared, since the single conductance measurements of PFO pores immediately after formation indicated inner pore diameters of 28.5 ± 5.6 nm, corresponding roughly to 35 to 55 PFO monomers. Assemblies with so many monomers typically form complete circles. The absence of significant changes in the conductance of PFO pores directly after their single-step formation of an open pore suggests that, in most cases, a stable circular assembly formed before the formation of an open pore, or a rearrangement to a closed circular assembly occurred during or immediately after forming an open pore. **Figure 1D** shows that the pores remained stable over the applied potential range from +100 mV to −100 mV, further supporting the formation of complete circular pore assemblies. Most experiments ended approximately 15 min after the pore formation due to spontaneous rupture of the lipid bilayer. We typically observed 3-5 PFO pore formations per hour in bilayers containing 30 mol% cholesterol, whereas no pore formation was observed in cholesterol-free planar lipid bilayers, as expected.

### Evaluation of PFO nanopores for estimating the volume and shape of analyte proteins

To test the performance of PFO nanopores for determining the volume and shape of single folded proteins, we used four test proteins: Fab fragment of polyclonal anti-biotin IgG (Fab, MW = 50 kDa), monoclonal anti-biotin immunoglobulin G (IgG, MW = 150 kDa,), β-Galactosidase (β-Gal, MW = 465 kDa, and Apoferritin (MW = 480 kDa). We added analyte proteins at a final concentration of ∼ 30 μg/mL to the *cis* side of the pore while applying a constant potential difference of 100 mV to the *trans* compartment of the lipid bilayer setup (**Figure 2A**). Resistive pulse measurements in the current recordings indicated that all tested proteins were able to enter PFO nanopores (**Figure 2C**, **Supplementary Figures S5** and **S6**), while control experiments revealed the absence of resistive pulses in the absence of analyte proteins, as expected. To achieve the largest frequency of resistive pulses, we polarized the bottom electrode in the *trans* compartment positively for experiments with the three test proteins and negatively for IgG. Analysis of the dwell time of resistive pulses *t_d_* revealed that the most probable dwell times for Fab, IgG, β-Gal, and Apoferritin were 285 μs, 265 μs, 120 μs, and 217 μs, respectively (**Supplementary Figure S7**). These residence times exceeded the diffusion time of a protein for the length of a PFO pore by two orders of magnitude. Such long residence times are often observed in biological nanopores, including those with large diameters, and are not fully understood at present. Possible reasons for extended residence times include electrophoretic forces, interactions between analytes and the nanopore walls, or complex flow patterns, such as vortices within the pore.^26,27,41^ The significant advantage of these extended residence times is that analyte molecules have the time to approach a representative sampling of all possible orientations within the pore, facilitating accurate estimation of protein volume and shape according to the theory developed by Golibersuch^42^ and others^21^.

**Figure 2.**
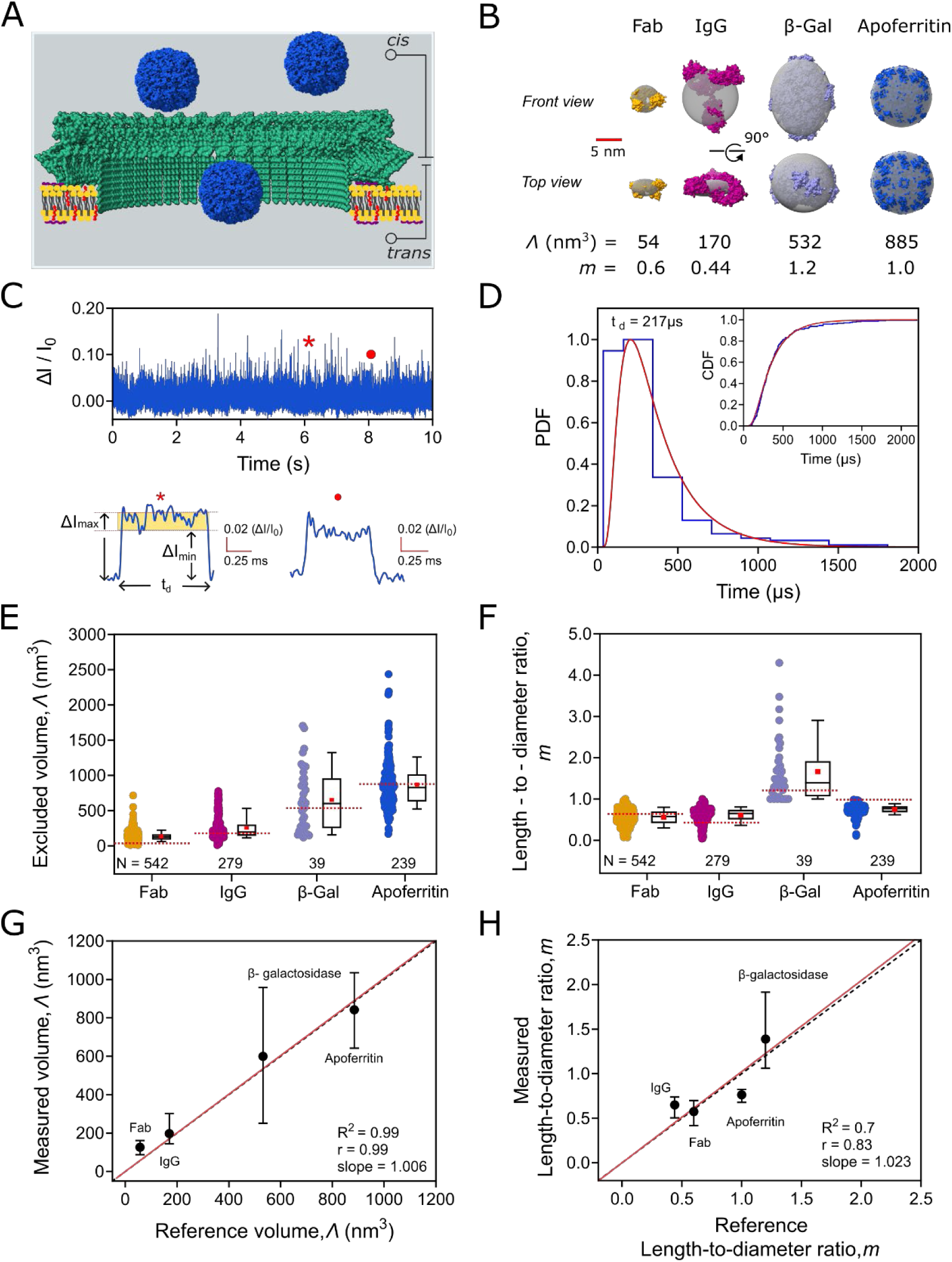
Estimation of the volume and shape (*i.e*., length-to-diameter ratio, *m*) of natively folded proteins based on resistive pulses recorded with PFO nanopores. (**A**) Schematic of the recording setup for single-molecule analysis using PFO pores embedded in a planar lipid bilayer. Schematic cross-sectional view of a PFO nanopore (green), along with a representative protein analyte (Apoferritin, blue) inside the pore. (**B**) Atomic structures of four test proteins and their respective reference ellipsoids of revolution (in grey) used to estimate their expected volume *Λ* and shape *m*. Fab fragment (Fab, 7FAB, MW 50 kDa, pI ≍ 8.7)^26^ in orange, anti-biotin immunoglobulin G (IgG,1HZH, MW 150 kDa, net charge at pH 7.5: *z* ≈ -4.2 × e)^22^ in magenta, β-Galactosidase (β-Gal, 1F4H, MW 465 kDa, pI ≈ 4.6) in purple, Apoferritin (6PXM, MW 480 kDa, pI ≈ 5) in blue. (**C**) Baseline–corrected current recording in the presence of Apoferritin showing resistive pulses as upward spikes. Recording buffer: 500 mM NaCl, 50 mM Tris-HCl, pH 7.5, 0.2 µM Amphipol, voltage = +100 mV applied to the bottom electrode in the *trans* compartment. Note: for recordings of Fab, β-Galactosidase and Apoferritin, we applied +100 mV to the bottom electrode in the *trans* compartment, and −100mV for IgG. The current recordings were collected at a 200 kHz sampling rate and filtered with a Gaussian low-pass filter at 10 kHz for Fab and 20 kHz for all other three proteins. Examples of two individual resistive pulses as marked in panel C. Maximum current blockage (Δ*I_max_*), minimum current blockage (Δ*I_min_*), and dwell time (*t_d_*) are illustrated. (**D**) Distribution of dwell times *t_d_* of resistive pulses from Apoferritin. The distribution includes the *t_d_* values of all detected resistive pulses longer than 20 μs and shows that the most probable *t_d_* value was 217 μs. The inset shows the cumulative density function of experimentally measured dwell times; we used this cumulative distribution to determine the most probable value of *t_d_* from a fit to all data, as this approach does not require binning. **(E**) Distribution of excluded volumes (*Λ*) and (**F**) length-to-diameter ratios (*m*) obtained from individual resistive pulses of four different proteins. Red horizontal dotted lines represent expected reference values from ellipsoid models of each protein. Horizontal lines in the boxplots represent median values. Solid red squares represent the experimental mean values. The box range represents the 25^th^ to 75^th^ percentiles, and the whisker range represents the 10^th^ to 90^th^ percentiles. (**G, H**) Plot of the median values of the excluded volume (*Λ*) and the length-to-diameter ratio (*m*) determined from single-event analyses of five proteins versus the expected reference values. Error bars show the first and third quartiles. The black dotted lines represent the ideal 1:1 agreement (slope = 1), and the red solid lines are the linear regressions performed imposing a zero intercept.

The amplitude of a resistive pulse contains information about the excluded volume *Λ* of an analyte particle, while current modulations during the resistive pulse are related to rotations of non-spherical particles in the electric field of a nanopore.^42^ Here, we approximate particle shape by the length-to-diameter ratio *m* of an ellipsoid of revolution with axes A, B, B and *m* = A/B with *m* < 1 corresponding to an oblate and *m* > 1 to a prolate.^7^ Based on this approach, we used a previously introduced data analysis package ^26^ to determine *Λ* and *m* from individual resistive pulses (**Supplementary Note 2**). To verify the experimentally determined parameters, we estimated reference values of *Λ* and *m* from available PDB structures of the tested proteins. In addition, we estimated reference values for protein volumes based on their molecular weights (**Figure 2B**; see **Supplementary Table S1**).

To obtain quantitative estimates of protein volume and shape from resistive pulse recordings, the nanopore lumen should be cylindrical.^21^ In addition, the pore diameter and length must be known. Since the measured open-pore current depends on both parameters, it is important to determine the pore length as precisely as possible to update the diameter at all times during the recording based on the measured open-pore current between resistive pulses. Here, we take advantage of the fact that multimeric biological nanopores such as poly C9, PLY, and PFO are typically composed of assemblies of the same protein, and these proteins are oriented in the pore in a fashion that does not change with the number of monomers in the pore assembly; we therefore assume pore length to be independent of pore diameter and constant. Observations of these pores by cryoEM or AFM support this assumption.^40,43,44^ Based on this assumption, we empirically determined the length of PFO pores using the four proteins in **Figure 2**. To do so, we plotted the experimentally measured excluded volumes of Fab, IgG, β-Gal, and Apoferritin against their known reference volumes, assuming pore lengths ranging from 7 to 11 nm to determine their volumes from the experimentally recorded blockade-current magnitudes (**Supplementary Figure S8**). We used a cutoff frequency of 10 kHz for Fab and 20 kHz for the Gaussian low-pass filter for all other test proteins to resolve resistive pulses from the noise and restricted the analysis of the shape and volume of these proteins to resistive pulses lasting at least 100 µs. A pore length of 9.8 nm provided the best agreement with all four reference volumes, yielding a slope of 1.006 ± 0.06 and a regression coefficient of *r* = 0.99 (**Supplementary Figure S8**). **Figure 2E-H** shows that, using a pore length of 9.8 nm, the analysis algorithm yields accurate median estimates of excluded volumes and approximate ellipsoidal shapes for all tested proteins. Importantly, accurate determination of the volume and shape of analyte proteins did not require a constant pore diameter from experiment to experiment or even during the same experiment, because the continuously measured open pore current between resistive pulses makes it possible to update the pore diameters for the analysis of each resistive pulse. With the length and diameter of PFO pores known at all times during a recording, volume and shape values from each resistive pulse can be determined.^21,26^ Importantly, this analysis does not require any calibration as long as the length of the pore is constant and the conductivity of the recording buffer is known, such that the pore diameter can be determined from the open pore current. This means that a new PFO pore, generated by a different operator in a different laboratory, directly reveals volume and shape measurements from recorded resistive pulses without calibration.

We attribute the broad distributions observed in both excluded volume and length-to-diameter ratios in **Figure 2E, F**, at least in part, to off-axis translocation within the pore.^45,46^ We demonstrated in previous work that analytes located away from the central axis inside a nanopore generate position- and orientation-dependent distortions in the recorded blockade signals.^47^ The effect of off-axis diffusion on the magnitude of resistive pulses increases as the diameter of the nanopore increases relative to the size of the analyte. We previously reported a similar broadening of the measured excluded-volume and shape distributions in which off-axis translocation produced asymmetric perturbations of the electric field within cylindrical nanopores, resulting in larger-than-expected resistive pulses.^7^ This effect is likely amplified in PFO pores due to their large diameter (∼ 30 nm) and relatively short length (9.8 nm), which permit substantial variation in analyte position and orientation during translocation. This effect will increase the fraction of volume estimates that exceed the reference value, and the proportion of shape estimates that are more extreme (i.e., either more pronounced oblate or more pronounced prolate shapes) than expected. Another factor that contributes to uncertainty in the determined volumes and shapes of large analyte proteins is that their longest dimension may exceed the length of the PFO nanopore. In this case, a lengthwise orientation of the analyte relative to the pore axis has the consequence that during the translocation, a part of the analyte particle is located outside of the zone of the strongest electric field in the centre of the nanopore. The contribution of these peripheral parts of the analyte particle to the amplitude of the resistive pulse will be smaller than assumed by the formula for the determination of volume. This effect will increase the fraction of volume estimates below the expected reference values and will be most pronounced for large molecules, as shown in the distributions for β-Gal and Apoferritin in **Figure 2E**. In terms of shape estimates, this effect will further increase the proportion of more extreme values (*i.e.*, either more pronounced oblate or more pronounced prolate shapes) than the expected reference values. The significantly steeper slope than 1.0 observed in **Figure 2H** may be due to this effect. Another effect that could contribute to broadening the distributions of determined protein volumes and shapes is deviation from a cylindrical shape of PFO pores. We propose, however, that the effect of ellipsoidal cross-sections of PFO nanopores on the amplitude modulations of resistive pulses is small since it has only a small effect on the resistance of a nanopore or on its access resistance as long as the ratio between major and minor axes is not significantly larger than 2 or 3. In addition, the electric field in an open pore with an ellipsoidal cross-section is expected to be homogeneous across the elliptical cross-section. While the presence of a particle may distort the electric field inside a pore with an elliptical cross-section more than in a circular cross-section, for most cases, these effects are likely minor unless the deviations from a circular cross-section become large. We exclude the possibility of large deviations from circular cross-sections based on the results shown below for the translocation of virus particles with a diameter of ∼ 20 nm. These particles can only translocate through a pore with an elliptical cross-section if the minor axis has at least the length of the particle diameter. For an elliptical pore with the same cross-sectional area as a cylindrical PFO pore with a diameter of 30 nm, elliptical cross-sections with an axis ratio of 2:1 have a minor axis with a length of 21.2 nm; axis ratios larger than 2:1 would result in a minor axis that is too short to facilitate the translocation of 20 nm particles; for instance, for an axis ratio of 3:1, the length of the minor axis is 17.3 nm.

### Application of PFO nanopores for characterizing rAAV-2 virus-like particles

To demonstrate application of PFO nanopores for characterization of large nanoparticles, we characterised empty recombinant adeno-associated virus type 2-like particles (rAAV2 VLPs) at the single-particle level. Recombinant AAVs are among the most versatile and clinically promising gene-delivery vectors, and a growing number of approved therapies and late-stage clinical trials rely on them.^50–52^ Their safety and efficacy depend on capsid integrity, accurate genome packaging, and the absence of defective or contaminating particles.^53,54^ However, rAAV preparations are heterogeneous and may contain full, empty, and partially filled capsids, as well as fragmented or aggregated species.^55,56^ While techniques such as transmission electron microscopy (TEM),^57^ mass photometry (MP),^57^ charge-detection mass spectrometry (CD-MS),^58^ and analytical ultracentrifugation (AUC)^59^ provide complementary information on rAAV structure and composition, the latter three of these techniques are ensemble-based and average across many particles; these techniques may therefore obscure rare or structurally distinct subpopulations. The resulting analytical need has motivated high-resolution, single-particle approaches, including label-free nanopore sensing in solution.^55,56^

The empty rAAV2 particles have a T = 1 icosahedral capsid with a reported diameter of approximately 24-26 nm (**Figure 3A**).^60^ Each capsid contains 60 viral proteins (VPs), typically represented as five copies of VP1, five copies of VP2, and 50 copies of VP3. ^61^ Reported molecular masses span approximately 81–87 kDa for VP1, 66–73 kDa for VP2, and 59–62 kDa for VP3.^61–63^ These values give a total capsid-protein molecular weight in the range of 3685 - 3900 kDa and, using a mass-to-volume conversion (see **Supplementary Note 4**), a protein-shell volume of 4437-4697 nm³ (mean 4567 nm³). For comparison, a sphere with a diameter of 24-26 nm has a total volume of 7238–9203 nm³ (mean 8221 nm³). Subtracting the estimated volume of the protein capsid shell from the total sphere volume gives an aqueous-core volume of 2541-4766 nm³ (mean, 3654 nm³), corresponding to a mean core diameter of approximately 19.0 nm and a simplified uniform-shell thickness of approximately 3.0 nm. DLS measurements in the nanopore recording buffer yielded an average hydrodynamic diameter of 21.8 ± 10.2 nm (**Supplementary Figure S9**), which is close to the reported 24–26 nm capsid diameter. The somewhat smaller average may reflect the high ionic strength of the recording buffer, whereas the broad uncertainty is consistent with the inability of ensemble-based DLS measurements to resolve particle-to-particle heterogeneity.

**Figure 3.**
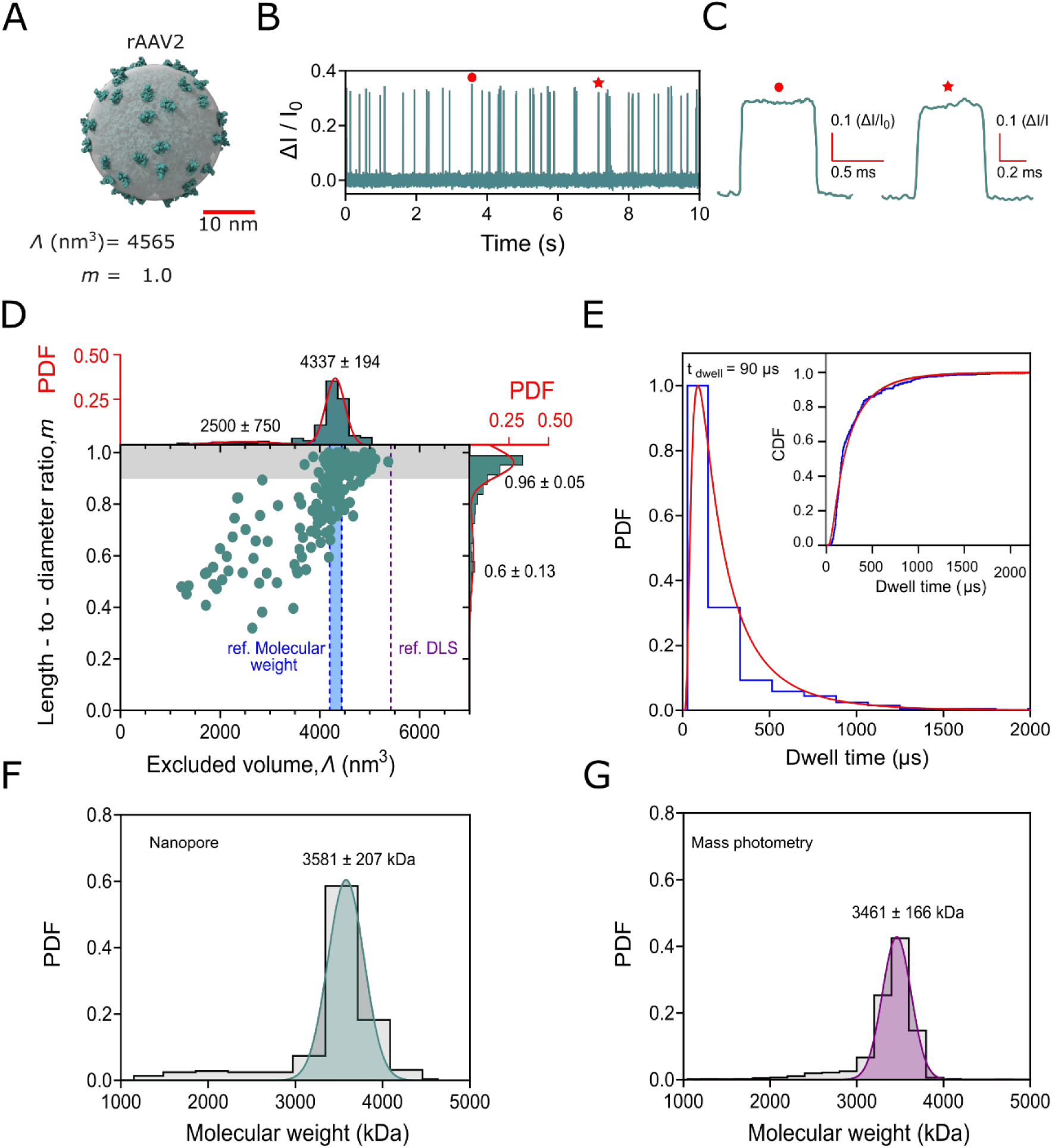
Characterization of empty recombinant adeno-associated virus type 2-like (rAAV2) particles with PFO nanopores. (**A**) Illustration of empty rAAV2 particles in green (PDB: 8FZ0) and corresponding spherical model with a diameter of ∼26 nm in transparent grey. (**B**) Baseline–corrected current recording through a PFO pore in the presence of rAAV2 showing resistive pulses as upward spikes. Recording buffer: 500 mM NaCl, 50 mM Tris-HCl, pH 7.5, 0.2 µM Amphipol, voltage = −100 mV applied to the bottom electrode in the *trans* compartment. The pI of empty rAAV2 particles is 6.3^48^; therefore, at pH 7.5, they are negatively charged and driven through PFO pores by EOF. Current recordings were collected with a 200 kHz sampling rate and filtered with a 20 kHz Gaussian low-pass filter for clarity. (**C**) Examples of two individual resistive pulses as marked in panel B. (**D**) Scatterplot and distributions of excluded volumes (*Λ*) and length-to-diameter ratio (*m*) obtained from individual resistive pulses of empty rAAV2 particles. The magenta-colored dotted horizontal line represents the expected reference volume of the complete particle, including its empty core, as determined by DLS measurements from our preparation, assuming a shell model for DLS analysis (see **Supplementary Figure S10**). The range of reference volumes shaded in blue was derived from converting the cumulative molecular weight of all 60 capsid proteins into the cumulative volume of the capsid shell (see **Supplementary Note 4**); since the molecular weights and, hence, molecular volumes of the three proteins in the viral capsid vary between sources, the area shaded in blue envelops the minimum and maximum cumulative volume. The range of *m* values shaded in grey indicates the virus particle population with *m* values of at least 0.9, *i.e*., particles are close to perfect spheres within the uncertainty of shape determination of at least *m* ± 0.1. (**E**) Distribution of dwell times (*t_d_*) of resistive pulses from rAAV2 particles. The distribution includes the *t_d_* values of all detected resistive pulses longer than 20 μs and shows that the most probable *t_d_* value is 90 μs. The inset shows the cumulative density function (CDF) of all experimentally measured dwell times; we used a fit to this CDF (red curve) to determine the most probable *t_d_* value. (**F**) Molecular weight (kDa) distributions of empty rAAV2 as determined by resistive pulses from PFO nanopores. We converted the estimated excluded volumes (*Λ*, nm^3^) to molecular weight using a volume-based molecular weight calculation (see **Supplementary Note 4**).^49^ (**G**) Molecular weight (kDa) distributions of empty rAAV2 particles as determined by mass photometry in the same buffer as used for nanopore recordings.

We only recorded resistive pulses from empty rAAV2 particles when -100 mV was applied to the electrode in the *trans* compartment (**Figure 3B**). Empty rAAV capsids have a reported isoelectric point of approximately pI = 6.3 and are therefore negatively charged at pH 7.5.^64^ Their motion toward the negatively polarised electrode suggests that translocation through the PFO pore is dominated by electroosmotic flow (EOF), rather than electrophoretic force (EPF). This condition is reasonable as PFO pores are structurally closely related to PLY pores (∼ 48% sequence identity)^65^ and the lumen of PLY pores is predominantly negatively charged at pH 7.5.^27^ We observed that the most probable dwell time for AAV2 particles is 90 μs (**Figure 3E**). This relatively long dwell time may result from weak interactions between rAAV2 particles and the luminal surface of PFO pores. The presence of these effects could slow down rAAV2 translocation, resulting in a dwell time being substantially longer than the estimated 1-D diffusion time of approximately 3 μs across the approximately 10 nm pore length. The capsid, which has a diameter of approximately 26 nm, is also expected to undergo rotational diffusion within the pore, where its orientation presumably randomizes within approximately 16 μs (corresponding to 3*τ_r_*, with *τ_r_* representing the rotational correlation time). Therefore, the observed dwell time of 90 μs enables sampling of most particle orientations as they move through the pores, which is required for the simultaneous estimation of volume and shape. Analyses of representative individual resistive pulses revealed that approximately 60% had uniform peak amplitudes (**Figure 3B**) and exhibited an almost complete absence of detectable intra-event current modulations beyond the baseline noise (**Figure 3C**). Withing the precision of shape determination, these results demonstrate that this fraction of virus particles has perfectly spherical shapes.

The excluded-volume distribution in **Figure 3D** shows two populations. The dominant population, accounting for approximately 75% of detected particles, is centered at *Λ* = 4337±194 nm³ (see **Supplementary figure S10** for rAAV2 for the estimation of empty rAAV2 VLP radius). This value is close to the theoretical volume estimate for the protein shell, 4437-4697 nm³, but markedly below the total particle volume, 7238-9203 nm³. We attribute this observation to the capsid architecture, which contains 12 symmetry-related pores, one at each fivefold vertex. Each pore forms a funnel-shaped channel with a minimum diameter of approximately 1.2 nm that widens to approximately 2.2 nm at the exterior. These channels are expected to admit water and small ions, consistent with direct observations of fluorophore penetration into the AAV capsid interior via the fivefold pore. These pores render the hollow core ionically connected to the surrounding electrolyte within an empty capsid.^17,66^ The resistive pulses in **Figure 3**, therefore, report predominantly the volume of electrolyte displaced by capsid protein rather than the electrolyte-filled core. Thus, under these conditions, rAAV2 behaves as a porous protein shell with an ionically conducting electrolyte-filled interior rather than as a sealed dielectric particle, although restricted ion transport and counterion accumulation may affect the effective conductivity of the interior compared to that of the bulk solution.

Mass photometry measurements shown in **Figure 3G** quantitatively supported the shell-only interpretation. These measurements determined a capsid mass at 3461±166 kDa, and the nanopore-derived molecular mass distribution agreed with this value within 4% (**Figure 3F**); both values also lie close to the 3685-3900 kDa range we calculated theoretically from nominal VP stoichiometry and reported VP masses, and they are consistent with previously reported mass photometry measurements of empty rAAV.^67,68^ Assuming an outer diameter of 25 nm as the average of the previously reported rAAV2 diameters, the most probable volume of the capsid from nanopore recordings of 4337 nm³ reveals a simplified uniform protein-shell thickness of 2.78 nm. Independently, the composition-based mean shell volume of 4567 nm³ gives a thickness of 2.98 nm. Both estimates are close to the approximately 2.6 nm radial span determined from cryo-EM density of empty AAV2 capsids.^69^ Because the capsid diameter approaches that of the PFO nanopore, we included the finite-size correction S (*d_m_*/D) in the calculation of the particle volume to account for the nonuniform current distribution around a large analyte (see equation 1),^21^

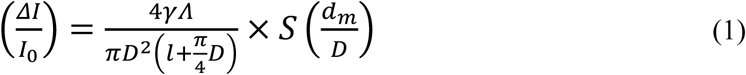

With *ΔI (*A) is the magnitude of the change in the current during translocation of a particle, *I*_0_ (A) is the baseline current, *γ* (unitless) is the electrical shape factor, *Λ* (m^3^) is the excluded volume of the particle, *D* (m) is the diameter of the cylindrical pore, *l* (m) is the length of the pore, 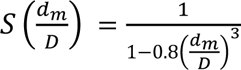 is the correction factor that depends on the relative values of *D* and the diameter of the molecule, *d_m_*.

Plotting the length-to-diameter ratio, *m*, against the excluded volume for individual resistive pulses (**Figure 3D**) further demonstrates that ∼58% of particles exhibited *m* ≥ 0.9, corresponding to a tightly clustered, high-volume population consistent with fully assembled, spherical capsids and representing the largest population in solution. A smaller subset retained volumes near that of a complete capsid, defined as *Λ* > 4143 nm³, one standard deviation below the center of the dominant peak, but had *m* < 0.9. These particles presumably represent incompletely closed or otherwise defective capsids; transient electro-deformation may also contribute because empty AAV capsids are shown to be more mechanically compliant than genome-containing capsids.^11^

The second population is smaller and broader, with a center at *Λ* = 2500 ± 750 nm³. All particles assigned to this population had *m* < 0.9, with *m* values centered at 0.60 ± 0.13, indicating oblate shapes. We attribute these remaining particles to partially assembled capsid fragments originating either from incomplete assembly or from dissociation of capsid fragments from the icosahedral capsid shell. In an independent geometric analysis, optimally fitted equal-volume ellipsoids of revolution yielded expected *m* values of ∼ 0.42 for half-shell fragments and ∼ 0.50 for three-quarter-shell fragments (see **Supplementary figure S11**), which agree well with the experimentally observed range. Although shape alone may not prove fragmentation, the progressive decrease of *m* with a decrease of *Λ* in **Figure 3D** provides evidence for the presence of incomplete capsids. The results presented here show that PFO nanopores can characterize rAAV2 capsids and capsid fragments at the single-particle level in terms of capsid-protein volume, approximate shape, and degree of assembly. The agreement between molecular-mass and DLS measurements supports quantitative volume analysis of the capsid shell, while the scatter plot of volume versus shape can resolve intact, near-spherical capsids from smaller, incomplete particles that may be hidden in population-based measurements. PFO nanopores, therefore, provide a robust, label-free method for assessing structural heterogeneity of viral particles.

### Single-particle characterization of ribosomes and their subunits

**Figure 4** demonstrates the capability of PFO nanopores to detect and characterize intact 70S ribosomes and their individual subunits at the single-particle level. We used structural models of the 30S subunit, 50S subunit, and 70S ribosome, all derived from the same PDB entry (7K00), to calculate theoretical excluded volumes and length-to-diameter ratios for each species (**Figure 4A**). **Figure 4B** shows that ribosome particles were able to enter the PFO nanopores after adding a solution containing 70S ribosomes to the *cis* chamber when applying a potential of −100 mV to the bottom electrode in the trans chamber. Representative individual resistive pulses of the ribosomal sample are shown in **Figure 4C** (see **Supplementary Figure S12**). Under the experimental conditions we used, the ribosome preparation contained intact 70S assemblies together with free 30S and 50S subunits, reflecting the dynamic equilibrium between association and dissociation of ribosomal components. The comparison of these nanopore-based observations with analysis of the same solution by mass photometry (**Figure 4D**) confirmed the presence of these three major species (see **Supplementary Figure S13** for detailed mass photometry data). Based on the areas of the three peaks, we determined the following relative abundances for the ribosome species: 45% for 30S, 26% for 50S, and 9% for 70S ribosomes by mass photometry. While the distribution of determined particle volumes from individual resistive pulses measured with PFO nanopores in **Figure 4E** revealed the following relative abundances: 43% for 30S, 19% for 50S, and 14% for 70S ribosomes. To assign resistive pulses to specific ribosome populations in **Figure 4E**, we fitted the cumulative distribution function (CDF) (see **Supplementary Note 6**) to the data, thereby avoiding binning artifacts associated with histogram-based methods. The CDF was fitted with a sum of three Gaussian components, initially constrained to a shared width of the 2^nd^ and 3^rd^ populations. The shared width was therefore retained in the final fit, yielding the mean (*μᵢ*) and common standard deviation (σ). These CDF-derived parameters were used to fit the PDF, with the mean and width, of each Gaussian component fixed accordingly. The data range for each population was defined as *μᵢ* ± σ, within which pulses were assigned to their corresponding species. The experimentally determined excluded volumes (*Λ*) showed good agreement with the theoretical values calculated from the corresponding ribosome structures (**Figure 4F**), demonstrating that PFO nanopores accurately capture the dimensions of individual ribosomal assemblies (30S, 50S, and 70S) from a heterogeneous mixture. The agreement with reference volumes for all three ribosomal particles confirms that resistive pulse amplitudes reflect the structural dimensions of individual particles and enables discrimination between ribosomal assembly states by their physical size. The PFO nanopores further revealed distinct morphological characteristics of each ribosomal species through the length-to-diameter ratio (*m*) (**Figure 4G**). The intact 70S ribosome exhibited a compact, oblate shape (*m* ≈ 0.8), consistent with the association of the 30S and 50S subunits into a globular ribonucleoprotein complex, whereas the isolated 30S and 50S subunits exhibited higher *m* values (∼1.6 and 1.2, respectively), reflecting their slightly elongated structures shown in **Figure 4A**. The consistency between nanopore-derived excluded volumes and length-to-diameter ratios across all analytes investigated in this study is further summarised in **Supplementary Figure S14** and demonstrates the overall performance of PFO nanopore-based measurements. These results show that PFO nanopores provide simultaneous access to particle dimensions and morphology of large biomolecular assemblies at the single-particle level.

**Figure 4.**
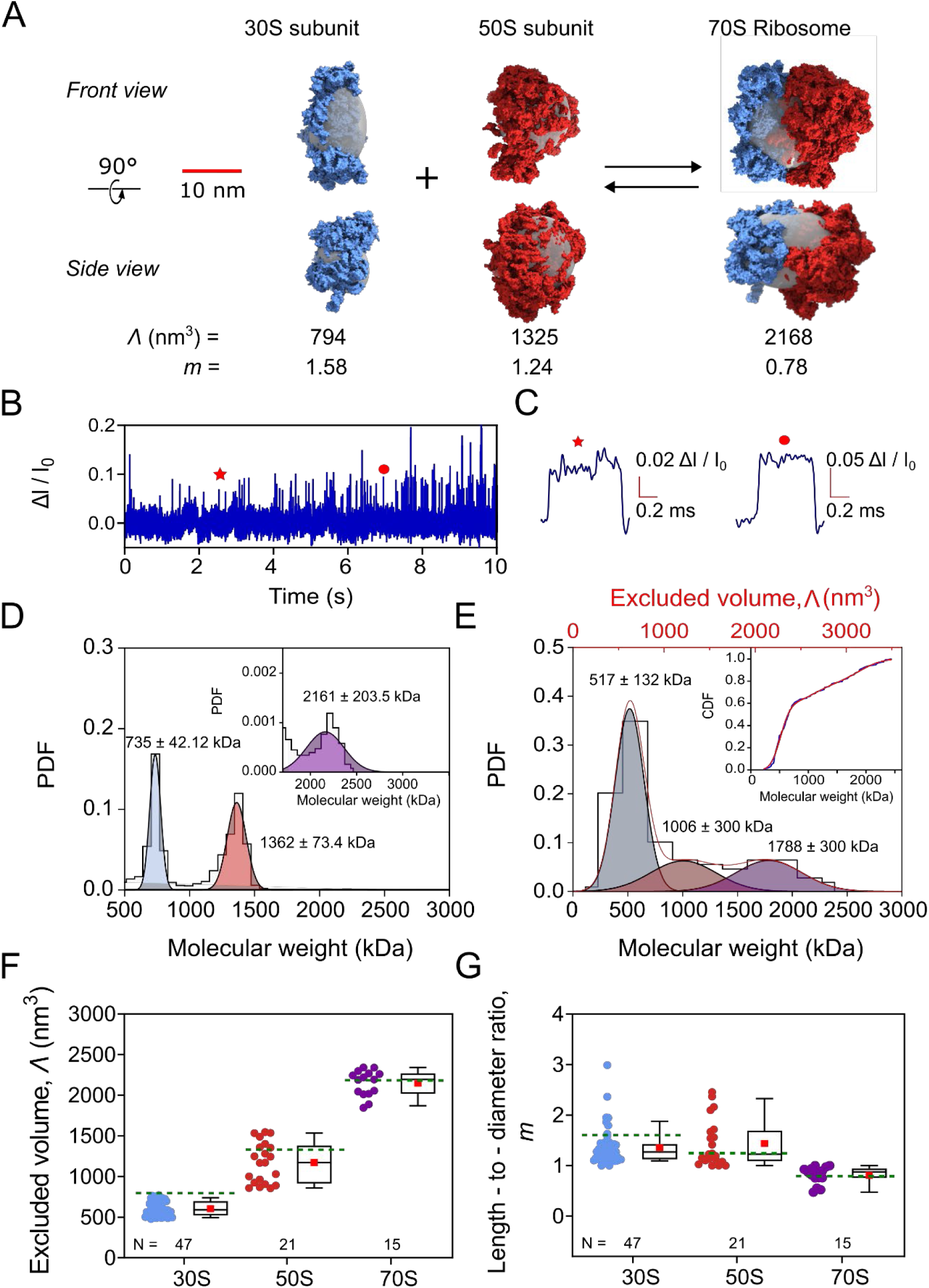
Characterization of the 70S ribosome and its subunits with PFO nanopores. (**A**) Schematic illustration of the 70S ribosome (PDB: 7K00) and its corresponding subunits: 50S and 30S. Structural models of the 30S, 50S, and 70S ribosomal particles were derived from the same PDB entry (7K00). The 30S and 50S subunits were extracted and analyzed separately to determine their theoretical volume and shape. (**B**) Baseline–corrected current recording in the presence of 70S ribosomes and their subunits showing resistive pulses as upward spikes. Recording buffer: 500 mM NaCl, 10 mM Mg(CH₃COO)₂, and 50 mM Tris-HCl at pH 7.5 with 0.2 µM Amphipol. A transmembrane voltage of −100 mV was applied to the bottom electrode in the trans compartment. Current traces were acquired with a 200 kHz sampling rate and filtered with a Gaussian low-pass filter with a cutoff frequency of 20 kHz for clarity. (**C**) Examples of two individual resistive pulses as marked in panel B. (**D**) Molecular weight distributions of the ribosome solution determined by mass photometry in the recording buffer. (**E**) Distribution of excluded volumes (*Λ*) obtained by nanopore recordings with PFO pores for the ribosome solution. The shaded areas result from a three-peak fit to the CDF shown in the inset (see **Supplementary Note 6** for details). (**F**) Excluded volumes ( *Λ*) and (**G**) length-to-diameter ratios (*m*) of individual resistive pulses, shown separately for each population. Events were assigned to a population if their *Λ* fell within ±1 SD of the corresponding peak center from the CDF fit in (**E**); events outside all three windows were excluded.

Together, these results establish PFO nanopores as a versatile platform for the label-free analysis of large, multicomponent biomolecular assemblies. By resolving intact ribosomal complexes and their individual subunits, PFO-based nanopore analysis provides direct access to the structural heterogeneity of large particles that is typically masked in ensemble measurements. The ability to simultaneously distinguish assembly state and particle shape highlights the potential of large-diameter biological nanopores for investigating complex biological systems at the single-particle level.

## Conclusions

This work introduces self-assembled Perfringolysin O (PFO) as a stable biological nanopore with an exceptionally large inner diameter of approximately 30 nm, enabling single-molecule characterization of biomolecules spanning a broad mass range from 50 kDa to 3.4 MDa. In addition to characterizing folded proteins, the large pore diameter enables direct resistive pulse analysis of viral particles and multicomponent ribonucleoprotein assemblies that were previously inaccessible to biological nanopores. The ability to resolve individual structural states, including complete viral capsids, capsid fragments, intact ribosomes, and their constituent subunits, demonstrates the versatility of PFO nanopores as a label-free platform for analyzing complex biological assemblies beyond molecular mass determination alone. Together, these findings establish large-diameter biological nanopores as powerful tools for structural fingerprinting of large biomolecules and provide a foundation for future applications in the characterization of complex biological assemblies.

## Materials and Methods

### Activation of PFO monomers and treatment with amphipol

PFO monomers (MyBioSource, MBS1166985), supplied at 0.7 mg/mL in 20 mM Tris–HCl, 0.5 M NaCl (pH 8.0) with glycerol, were first dialysed against 50 mM Tris–HCl buffer (pH 7.5) for 10 min using Slide-A-Lyzer MINI dialysis devices with a 10 kDa molecular weight cut-off (Thermo Fisher Scientific, 69570) to remove excess glycerol. We determined the protein concentration after dialysis by measuring absorbance at 280 nm using a NanoDrop and an extinction coefficient of 74,260 M⁻¹ cm⁻¹ ^70,71,^ resulting in a final concentration of 0.5 mg/mL. We aliquoted the protein into 10 µL volumes, flash-frozen them, and stored them at −80 °C. We activated PFO monomers (0.5 mg mL⁻¹, 10 µL) by the addition of 1.2 µL of a freshly prepared 70 mM 1,4-dithiothreitol (DTT; Sigma-Aldrich, 10197777001) solution in 50 mM Tris–HCl buffer (pH 7.5, Thermo Scientific) for 10 min at 37 °C, without shaking. We then incubated activated PFO monomers with amphipol A8-35 (Anatrace), maintaining a PFO: amphipol molar ratio of 12:1 in 50 mM Tris−HCl (pH 7.5) for 48 h at 37 ⁰C, without shaking. The final concentration of PFO monomers was set to 0.25 mg mL-1.

### Planar lipid bilayer formation and characterization

We prepared a lipid mixture containing 70 mol% 1,2-diphytanoyl-sn-glycero-3-phosphocholine (DiphyPC, Sigma-Aldrich/Merk 850356P) and 30 mol% cholesterol (Fisher Scientific 11493310) in n-octane (Sigma-Aldrich/Merk 296988) with a final concentration of 15 mg mL-1 DiphyPC. For current measurements with planar lipid bilayers, we used an integrated chip-based, sixteen-channel parallel bilayer recording setup (Orbit 16 TC, Nanion Technologies) with multielectrode cavity-array chips (MECA chips, Ionera) and EDR3 software (Elements). MECA chips bearing sixteen channels with a cavity diameter of 100 µm were used to form free-standing lipid bilayers according to the manufacturer’s instructions. Briefly, we added 150 µL of recording buffer (500 mM NaCl and 50 mM Tris–HCl, 0.2 µM amphipol, pH 7.5) to the cis compartment of the chip. After positioning the magnetic stir bar, we deposited 0.2 µL of lipid solution directly on the top and bottom of the chip’s surface in the stir bar’s trajectory. The stir bar was rotated several times to spread out the lipids. We confirmed the quality of the resulting lipid bilayers by verifying that the baseline current remained within 0 ± 0.3 nA while applying potential differences of ±100 mV, and that the electrical capacitance was 18 ± 5 pF. The recording software EDR3 automatically estimated the membrane capacitance by analysing the current response to an applied triangular potential difference. We allowed the bilayers to stabilise for 5 min. During this time, we tested the stability of the bilayers by applying transmembrane voltages of ±100 mV for 1 min and confirming the expected noise level, the absence of leak currents, and that the capacitance remained above a threshold of 15 pF.

### Assembly of a PFO nanopore in a planar lipid bilayer

We added 5 μL of incubated 0.25 mg/ml PFO solution with amphipol to the cis compartment of the MECA chip and maintained an applied potential difference of +100 mV until we observed a sudden single-step current jump, indicating successful formation of a PFO nanopore in the lipid bilayer. In a typical experiment, a nanopore formed within 10-15 min of adding the PFO-amphipol solution. We calculated the inner diameter of the pore and the corresponding number of PFO monomers from the pore conductance measured as a difference between baseline and open pore current. We estimated the pore diameter using the equation proposed by Cruickshank et al^72^, which assumes the pore to be perfectly cylindrical and accounts for both the resistance of the nanopore and the access resistances:

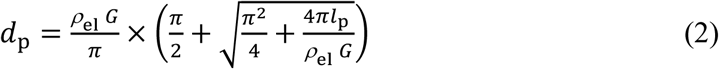

In equation 2, *d*_p_ [m] is the inner diameter of the cylindrical nanopore, *l*_p_ [m] is the pore length, ρ_el_ [Ω·m] is the electrical resistivity of the electrolyte buffer, and *G* [S] is the conductance of the pore. We determined a channel length of *l*_p_ = 9.8·10^-9^ m (**Supplementary Figure S8**). We then calculated the number of PFO monomers comprising the nanopore using a previously published geometric model. ^73^ We conducted all electrical measurements at a sampling rate of 200 kHz and a current range of 20 nA selected via the EDR3 software.

### Resistive pulse sensing experiments with protein analytes

After forming a PFO nanopore in the planar lipid bilayer, we performed a stepwise voltage sweep from −100 mV to +100 mV to verify pore stability and determine the offset current at an applied potential difference of 0 mV. We added the analyte protein solution to the cis compartment of the chip; the final protein concentration is specified in the corresponding section below. To drive translocation of analyte proteins through the nanopore, we applied a constant transmembrane potential of ±100 mV. Voltage sweeps were repeated every 5 min to monitor nanopore stability and to assess potential changes in the offset current. When an offset potential was detected, the recorded currents were corrected accordingly.

### Handling of analyte proteins

We used Fab protein (Fab Biotin Polyclonal Antibody, Thermo Scientific, cat. #800-101-098), anti-biotin immunoglobulin G (IgG, Sigma-Aldrich, B3640), β-Galactosidase (β-Gal, Sigma-Aldrich G3153), and Apoferritin (Sigma-Aldrich, A3660) in nanopore experiments. Fab and IgG were used as received, while β-galactosidase was resuspended in PBS buffer (ROTI Cell, 9143), pH 7.5, to prepare the stock solutions. β-Galactosidase and Apoferritin were purified and fractionated by size-exclusion high-performance liquid chromatography (HPLC). The final molar concentrations of analyte proteins in the cis compartment were 0.65 μM for FAB, 0.21 μM for IgG, 0.11 μM for β-Gal, and 0.09 μM for Apoferritin. Empty Adenovirus serotype 2 virus-like particles (AAV2 VLP) (Creative Diagnostics, DAG-WT824) were used at a concentration of 2.67 × 10¹¹ VLPs/mL. 70S ribosomes from *E.Coli* (New England Biolabs, P0763S) were used at a final concentration of 0.43 μM.

### Analysis of nanopore data

Unless otherwise noted, we filtered the acquired data with a Gaussian low-pass filter at a cut-off frequency of 10 kHz for Fab and 20 kHz for all other analytes and performed a threshold search (5 × the standard deviation of the baseline current) for the detection of resistive pulses.

### Mass Photometry

We acquired all mass photometry data using a TwoMP mass photometer (Refeyn Ltd., Oxford, UK), similar to that described previously by Kukura *et al.*^74,^ with slight modifications. Briefly, we cleaned microscope coverslips (24 × 50 mm, Thorlabs, cat. no. CG15KH1) with alternating washes of isopropanol and pure water, repeated three times. We used PDMS CultureWell gaskets (cat. no. GBL103250) to define compartments for mass analysis. Ribosomes were diluted to a final concentration of 26 nM in 50 mM Tris-HCl recording buffer containing 500 mM NaCl and 10 mM Mg(CH₃COO)₂, pH of 7.5, immediately before mass photometry measurements. For each mass photometry acquisition, 18 μL of buffer was first added to the chamber and autofocus was stabilised. Then, 2 μL of diluted protein was added, and movies lasting 180 s were recorded. For mass photometry data on AAV2 particles, we acquired the data as described by *Liang et al.* ^50^, with slight modifications.^90^ Briefly, 2 μL of 1.6 × 10^10^ VLP/ml of rAAV2 particles were added to 18 μL of buffer containing 500 mM NaCl, 50 mM Tris pH 7.5, after autofocus stabilisation, and movies of 180 s duration were recorded. Mass photometry data acquisition was performed using AcquireMP software (Refeyn Ltd., v2.4.1). Contrast-to-mass calibration was performed using MassFerence P1 Calibrant (Refeyn Ltd., MP-CON-41033).

## AUTHOR INFORMATION

A.V. acknowledges financial support from the Swiss National Science Foundation (Grant number: 200020_197239) and from the Adolphe Merkle Foundation. A. M. acknowledges financial support from the Swiss National Science Foundation (SNSF) “SPARK” funding (CRSK-2_221078), support from the SNSF through the National Center of Competence in Research (NCCR) Bio-Inspired Materials, Novartis Foundation for Medical-Biological Research, as well as the Adolphe Merkle Institute. M.M. acknowledges financial support from the Swiss National Science Foundation (Grant number: 200020_197239) and from the Adolphe Merkle Foundation.

