## Supplementary Information for "Characterization of Single Ribosomes and Virus Particles with Large-Diameter Perfringolysin O Nanopores"

***Supplementary Note 1: Estimation of inner pore diameter and number of monomers***

We calculated the equivalent PFO pore diameter based on its conductance by comparing the baseline current with the current after single-pore insertion, following the protocol reported by Chanakul *et al.*[1] for poly(C9) nanopores. We used the equation presented by Cruickshank *et al.*[1]

$$d_{\mathrm{p}} = \frac{\rho_{\mathrm{el}}\, G}{\pi} \times \left(\frac{\pi}{2} + \sqrt{\frac{\pi^2}{4} + \frac{4\pi l_{\mathrm{p}}}{\rho_{\mathrm{el}}\, G}}\right) \tag{1}$$

In this equation:

$d_{\mathrm{p}}$ (m) - Inner diameter of the nanopore,

$\rho_{\mathrm{el}}$ $(\Omega \cdot \mathrm{m})$ - Electrical resistivity of the electrolyte buffer. we used an experimentally measured value of conductivity of 4.23 $\mathrm{S/m}$ for the recording buffer with 500 mM NaCl,

$G$ $(\Omega^{-1})$ - Conductance, determined as the ratio of the difference between the baseline current and the open pore current at the applied voltage,

$l_{\mathrm{p}}$ (m) - Pore length was estimated to be 9.8 nm (see **Supplementary Figure S4**) and assumed to be constant for all pore diameters.

We then used the determined pore diameter to estimate the number of PFO monomers comprising the pore using a geometric model described by Fennouri *et al.*[2]

$$d_p = s\sqrt{\frac{1}{\pi}\left(\frac{n}{tan\left(\frac{\pi}{n}\right)} - \frac{(n-2)\pi}{2}\right)} \tag{2}$$

In this equation:

$d_{\mathrm{p}}$ (m) - Inner diameter of the nanopore, obtained from Equation (1),

$n$ - Number of monomers in the PFO pore,

$s$ (m) - Diameter of a cylindrical rod representing a PLY monomer.

The model assumes that the PFO pore is a circular arrangement of cylindrical rods (PFO monomers) oriented perpendicular to the surface of a planar lipid bilayer. These rods are positioned at the corner points of a regular polygon, with the diameter of each rod equal to the distance between the rods.

We have previously demonstrated[1,3,4] that a Taylor expansion of Equation (2) provides an accurate estimation of pore diameter:

$$d \approx s(0.318n - 0.784) \tag{3}$$

Since no cryo-EM pore structure is currently available for Perfringolysin O, we approximated the monomer dimensions using structural information from Pneumolysin (PLY). PLY and PFO are homologous cholesterol-dependent cytolysins with highly conserved domain architecture and comparable monomeric folds.[5]

Due to the strong structural similarity between PLY and PFO, we used the same monomer diameter to represent PFO in our geometric model. To determine the diameter $s$ of a cylindrical PFO monomer, we used Equation (3) and information from a recent PLY cryo-EM structure [6] where the inner diameter of PLY was found to be 22 nm, and the pore consisted of 42 monomers. Consequently, the diameter, s, of a cylindrical rod representing a PLY monomer in a PLY nanopore and therefore in PFO is approximately 2.02 nm.[4]

***Supplementary Note 2: Data analysis algorithm***

We developed data analysis software to obtain the excluded volume $\Lambda$ and length-to-diameter ratio $m$ of a target analyte from nanopore recordings.

The analysis consists of three sequential steps:

<u>1. Baseline search</u>. In the first step, the recording $x$, composed of $W$ Samples are imported and digitally filtered using a Gaussian low-pass filter with the desired cutoff frequency. The data are subsequently processed with a custom-made baseline search algorithm that operates as follows:

1) The filtered recording $x^{\mathrm{f}}$, with size $W$, is subdivided into $N+1$ segments, $N$ containing $M$ samples each and one containing the remaining $W-NM$ samples. The value of $M$ is set by the user and should always be M >100 and at least 30 times longer than the longest translocation event.
2) An empty array $x^{\mathrm{b}}$ with length $W$ is initialised, which will be filled with the estimated baseline values. For clarity, the $j$ th element of an array $x$ is indicated as $x[j]$.
3) The mean ($\bar{y}_{\mathrm{i}}$) and the standard deviation ($\sigma_{\mathrm{i}}$) of each segment $y_i$ is calculated.
4) The minimum standard deviation $\sigma_{\mathrm{min}}$is identified as the minimum value among the standard deviations of the $M$-sized segments $y_i$.
5) A threshold parameter $\tau > 1$ is defined (typically 1.2< $\tau$ <1.6).
6) For each segment $y_i$, with $i = 1 \dots N$, the standard deviation $\sigma_{\mathrm{i}}$ is compared to $\tau\sigma_{\mathrm{min}}$ with two possible outcomes:
   A) if $\sigma_{\mathrm{i}} \leq \tau\sigma_{\mathrm{min}}$, the elements of $x^{\mathrm{b}}$ comprised between $x^{\mathrm{b}}[(i-1)M]$ and $x^{\mathrm{b}}[iM]$ are filled with the average value $\bar{y}_{\mathrm{i}}$

B) if $\sigma_{\mathrm{i}} > \tau\sigma_{\mathrm{min}}$, $y_i$ is subdivided into 10 samples, $z_k^i$ ($k = 1 \dots 10$) of length M/10. The elements of $x^{\mathrm{b}}$ comprised between $x^{\mathrm{b}}[(i-1)M + M(k-1)/10]$ and $x^{\mathrm{b}}[(i-1)M + Mk/10]$ are filled with the average value of the $z_k^i$ segment, $\bar{z}_k^i$.

7) The baseline is finally completed by filling in the last $W - NM$ elements of $x^{\mathrm{b}}$ with the mean value of the last $W - NM$ elements of $x^{\mathrm{f}}$.

The baseline trace $x^{\mathrm{b}}$ is important because it allows tracking step changes in the baseline current and, hence, changes in the diameter of the nanopore over time.

2. Event detection. After determining the baseline trace, the normalised ($\Delta I/I_0$) trace, $x^{\mathrm{n}}$ is calculated as $x^{\mathrm{n}} = (x^{\mathrm{f}} - x^{\mathrm{b}})/x^{\mathrm{b}}$ and processed with the following threshold-based event detection algorithm:

1) The trace $x^{\mathrm{n}}$ is subdivided into $F + 1$ segments, $F$ containing $G$ sample each and one containing the remaining $W - FG$ samples.
2) For each $G$-sized segment $q_i$, with $i = 1 \dots F$, the mean $\bar{q}_i$ and the standard deviation $\sigma_i$ are computed. Two thresholds are calculated, $T_{\mathrm{s}} = \bar{q} + u\sigma_i$, where $u > 1$ (typically $u > 4$), and $T_{\mathrm{e}} = \bar{q} + \sigma_i$.
3) A preliminary search is performed to identify at which position in $q_i$ the signal first exceeds $T_{\mathrm{s}}$ ($e_{\mathrm{s}}$, event start) and then falls below $T_{\mathrm{e}}$ ($e_e$, event end). For each event, the end and start positions of the events in the trace $x^{\mathrm{n}}$ are calculated to include part of the baseline as $E_{\mathrm{s}} = e_{\mathrm{s}} + (i-1)G - 40$ and $E_{\mathrm{e}} = e_{\mathrm{e}} + (i-1)G + 40$, and stored (extended events).
4) The detected events are subsequently refined, as a simple threshold-based search could fragment a single translocation into multiple short events, or interpret noise spikes as translocation events:
   A) All samples in $x^{\mathrm{n}}$ with value smaller than $T_{\mathrm{s}}$ are stored in an array $w$ of size $R$.
   B) A matrix $B_{(R-Q)\times Q}$ is constructed by stacking Q delayed copies of $w$, such that the j$^{\mathrm{th}}$ column of $B$ contains the elements of $w$ comprised between $w[j]$ and $w[R - Q + j]$.
   C) The standard deviation of each row of $B$ is calculated, and the minimum value obtained is defined $\sigma_0$.
   D) Each extended event is fitted using two Gaussian peaks having the same standard deviation $\sigma_0$ and centred at positions $L_0$ and $L_1 (L_1 > L_0)$. $L_0$ and $L_1$ represent an estimate of the baseline and of the event amplitude, respectively.
   E) The event is fitted with a 2-level step-fitting algorithm using $L_0$ and $L_1$ as levels.
   F) The portion of the event which is best fitted by $L_1$ is considered as the refined event. If the duration of the event exceeds a user-defined minimum duration, the event is accepted and stored.

3. Event analysis. The refined resistive pulses are analysed to determine the minimum and the maximum intensity values ($dI_{\mathrm{min}}, dI_{\mathrm{max}}$), which are needed to determine the volume ($\Lambda$) and the ellipsoid-

equivalent shape ($m$) of the analyte particles. Since the measurements are affected by noise, the absolute maximum and minimum of the event trace are not good estimates of $dI_{\min}$ and $dI_{\max}$. Instead, we developed the following method:

1) A matrix $B_{(R-Q)\times Q}$ is constructed as described above. The 20th and the 80th percentiles of each row of $B$ are calculated and stored in two arrays, $P_{20}$ and $P_{80}$, with sizes $(R-Q)$.
2) The minimum value of the array obtained from the elementwise difference $P_{80} - P_{20}$ is labelled $dI_0$.
3) The 20th and the 80th percentile of each event, $dI_{20}$ and $dI_{80}$ is computed.
4) $dI_{\min}$ and $dI_{\max}$ are calculated as $dI_{\min} = dI_{20} + dI_0/2$ and $dI_{\max} = dI_{80} - dI_0/2$.
5) $dI_{\min}$ and $dI_{\max}$ are used to compute $\Lambda$ and $m$ as described in by Yusko *et al.*[7] and Houghtaling *et al.*[8]

***Supplementary Note 3: Approximation of protein shape with an ellipsoid of rotation***

Protein structures obtained from the PDB were converted to solvent-excluded surfaces using a custom algorithm. Protein volumes were then calculated based on these solvent-excluded surfaces. To estimate the ellipsoid parameters (a, a, b), the minimum volume enclosing ellipsoid (MVEE) algorithm was applied, which optimises the semiaxes based on the solvent-excluded surface to determine the best-fit ellipsoid shape.[9]

***Supplementary Note 4: Approximation of protein molecular weight from its radius***

We calculated the molecular weight of a single protein from its radius, assuming a spherical shape of the particles, using Equations 3 and 4.[10]

$$r = 0.066 * (M.W)^{1/3} \quad (3)$$

$$V = \frac{4}{3} * \pi * r^3 \quad (4)$$

Where,

$r$ is the radius of the protein, assuming a spherical model for its shape, nm.

$M.W.$ is the molecular weight of protein, Da.

$V$ is the volume-based molecular weight, nm$^3$.

### *Supplementary Note 5: DLS measurement of rAAV2 VLP*

Hydrodynamic size measurements of AAV2 virus-like particles (VLPs) were performed using an LS Spectrometer™ II (LS Instruments AG, Fribourg, Switzerland). Samples containing 2.67 × $10^{11}$ VLPs $mL^{-1}$ in 500 mM NaCl and 50 mM Tris-HCl (pH 7.5) were measured in cylindrical glass cuvettes at 25 °C. Scattered light was collected at a backscattering angle of 175° using a 660 nm laser. For each sample, 15 autocorrelation functions were acquired over 30 s and averaged. The hydrodynamic diameter was determined by fitting the averaged intensity autocorrelation function using the CONTIN algorithm.

### *Supplementary Note 6: Gaussian CDF Fitting for Discrimination of Ribosome populations*

To assign resistive pulses to specific ribosome populations in Figure 4E, we fitted the cumulative distribution function (CDF) thereby avoiding binning artifacts associated with histogram-based methods. The molecular weight cumulative distribution was fitted using a three-component Gaussian mixture model. The cumulative distribution function was represented as :

$$F(x) = \sum_{i=1}^{3} w_i \Phi\left(\frac{x - \mu_i}{\sigma_i}\right) \tag{5}$$

Where:

$w_i$ represents the fractional contribution of each population ($\sum_i w_i = 1$)

Φ is the standard normal cumulative distribution function

$\mu_i$ is the mean molecular weight

$\sigma_i$ is the standard deviation

*Supplementary Figures:*

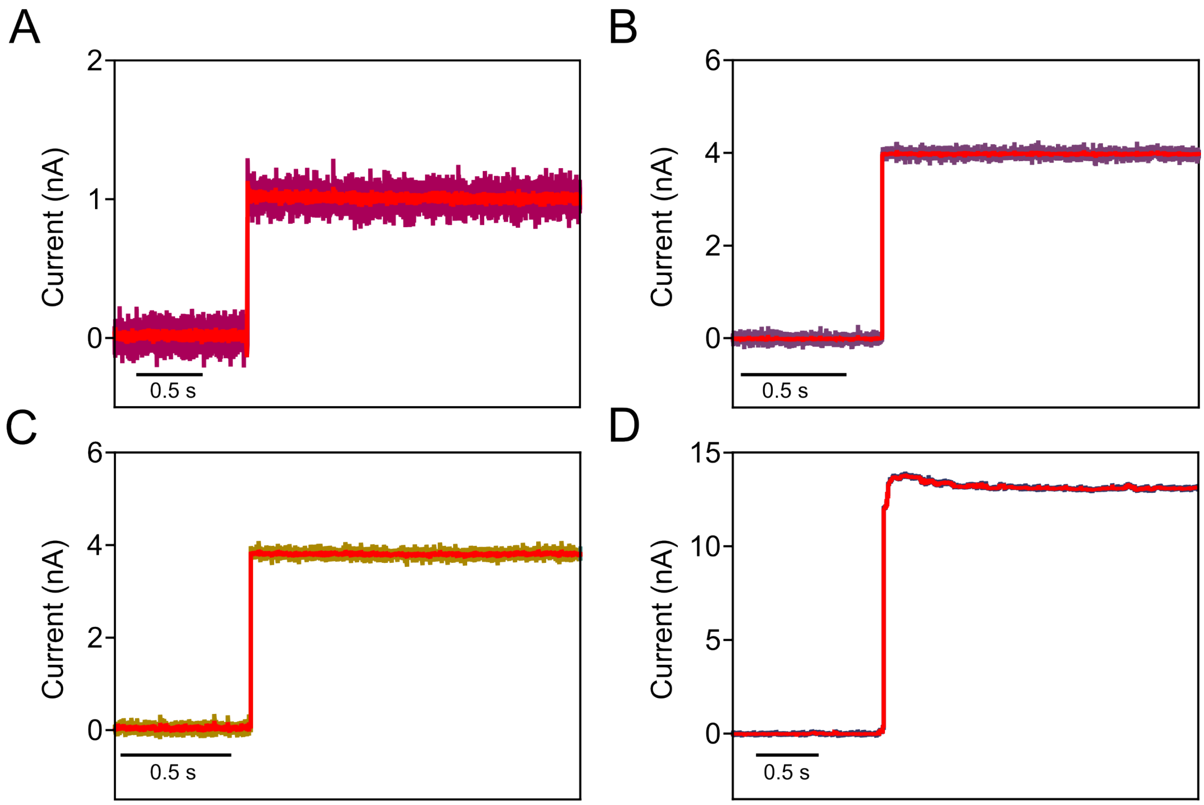

**Supplementary Figure S1. Single-pore insertions of PFO into lipid bilayers at +100 mV under varying ionic conditions.** Current recordings were obtained using a recording buffer (pH 7.5) containing 50 mM Tris-HCl and 0.2 µM amphipol, with NaCl concentrations of (**A**) 50 mM, (**B**) 150 mM, (**C**) 300 mM, and (**D**) 1 M. All recordings were collected at a sampling rate of 200 kHz. Traces shown in red represent the same data after additional low-pass filtering at 10 kHz. A 10 kHz cutoff frequency was chosen for display purposes only; details of the cutoff frequencies used for quantitative data analysis are provided in the Methods section.

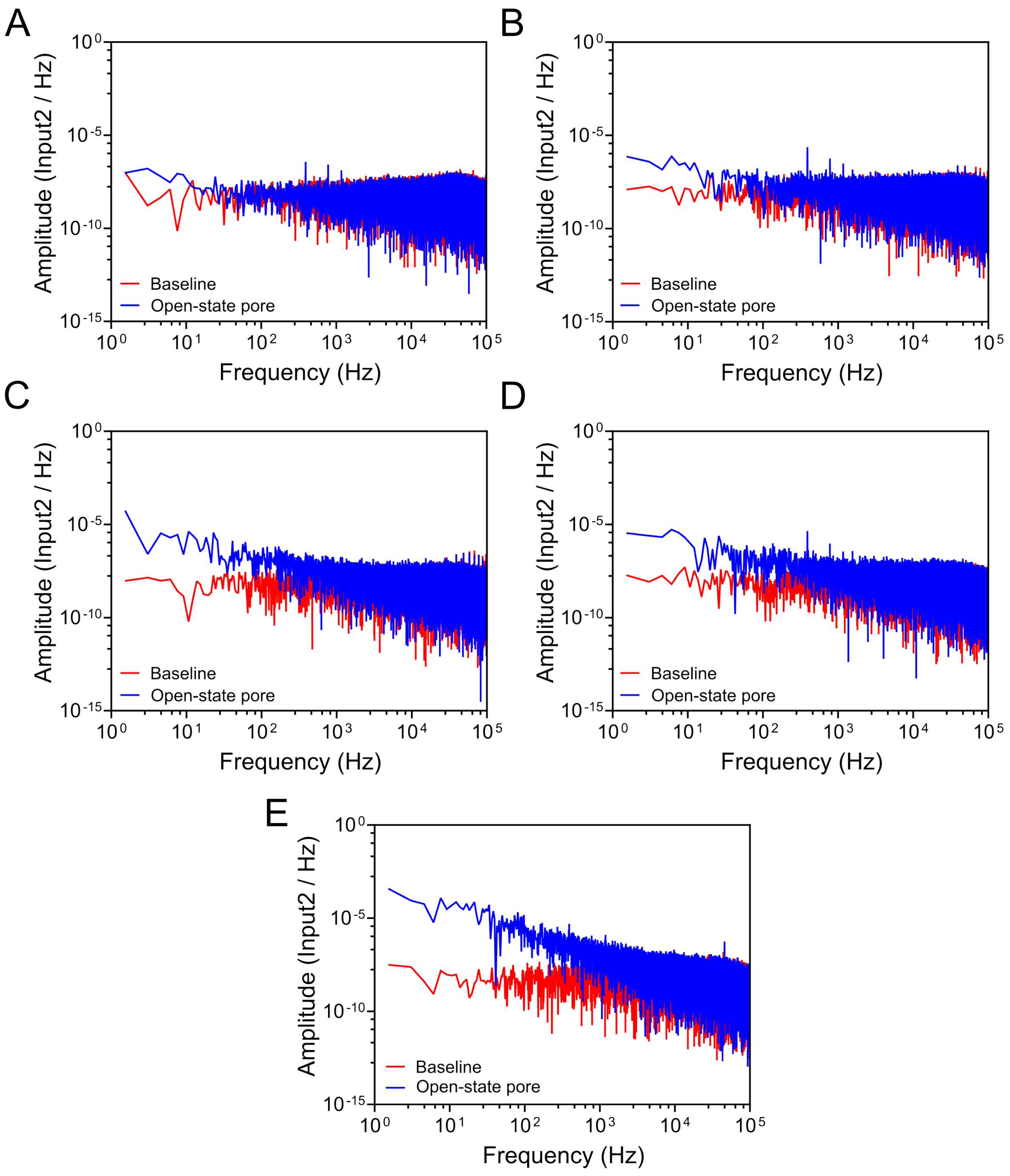

**Supplementary Figure S2. Effect of NaCl concentration in the recording buffer on the current noise from recording with a single PFO pore.** Comparison of power spectral densities (PSD) of the current before (baseline, red) and after insertion of a single PFO pore (open-state pore, blue) using a recording buffer (pH 7.5containing 50 mM Tris-HCl and 0.2 μM amphipol at NaCl concentrations of (**A**) 50 mM, (**B**) 150 mM, (**C**) 300 mM, (**D**) 500 mM, and (**E**) 1 M. The PSD was determined from the current recordings collected with a 200 kHz sampling rate without filtering.

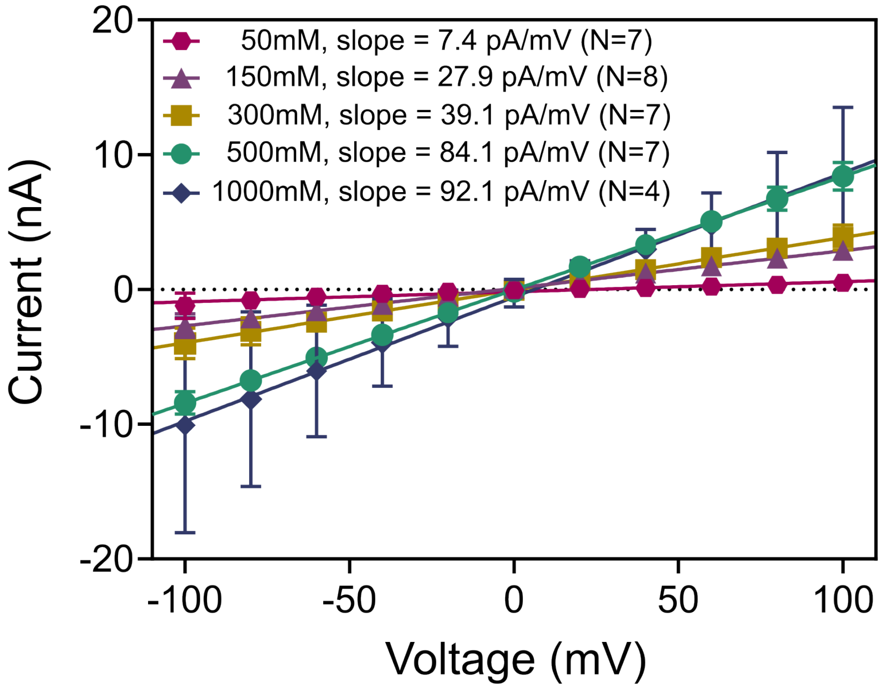

**Supplementary Figure S3. Current-voltage (I–V) curves of PFO pores in recording buffer at different ionic strengths.** The recording buffer, pH 7.5, contains 50 mM Tris-HCl, 0.2 µM amphipol, with 50 mM, 150 mM, 300 mM, 500 mM, and 1 M NaCl. Error bars represent the standard deviations calculated from a minimum of three repeats.

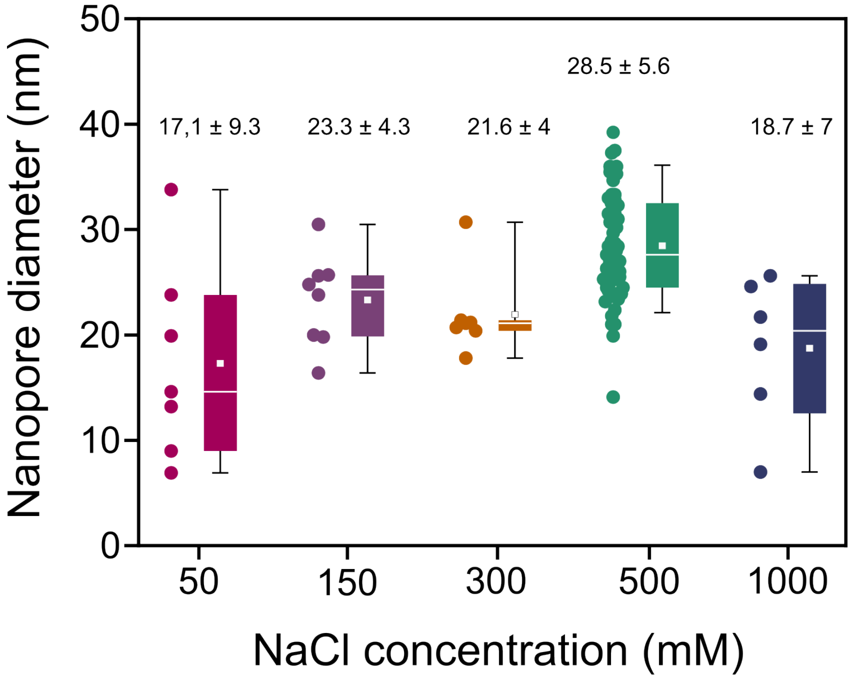

**Supplementary Figure S4. Estimates of the inner diameter of the PFO pore at different NaCl concentrations.** Calculations of the diameter employed a pore length of 9.8 nm, as determined from length calibration (see **Supplementary Figure S8**). Solid white squares indicate mean values, whereas median values are shown as white horizontal lines. The box range corresponds to the $25^{th}$ to $75^{th}$ percentile, whereas the whisker range corresponds to the $10^{th}$ to $90^{th}$ percentile.

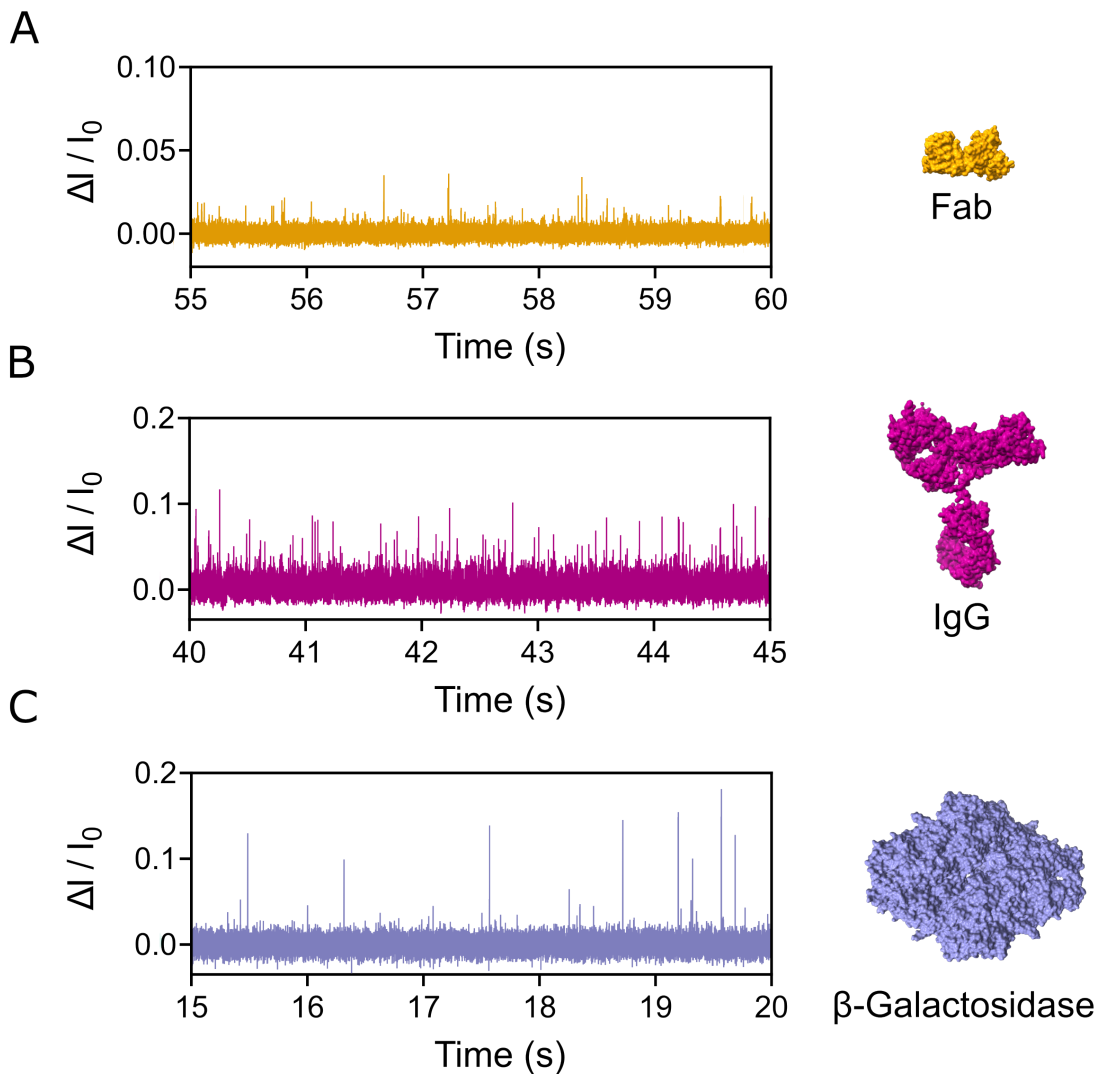

**Supplementary Figure S5. Baseline–corrected current recordings showing resistive pulses (upward spikes) measured with PFO nanopores** in the presence of (**A**) Fab, (**B**) IgG, (**C**) β-Galactosidase. The current recordings were collected in a buffer containing 500 mM NaCl, 50 mM Tris-HCl and 0.2 μM amphipol, pH 7.5 with a 200 kHz sampling rate and were filtered with a 10 kHz Gaussian low–pass filter for Fab and a 20 kHz Gaussian low–pass filter for IgG and β-Galactosidase.

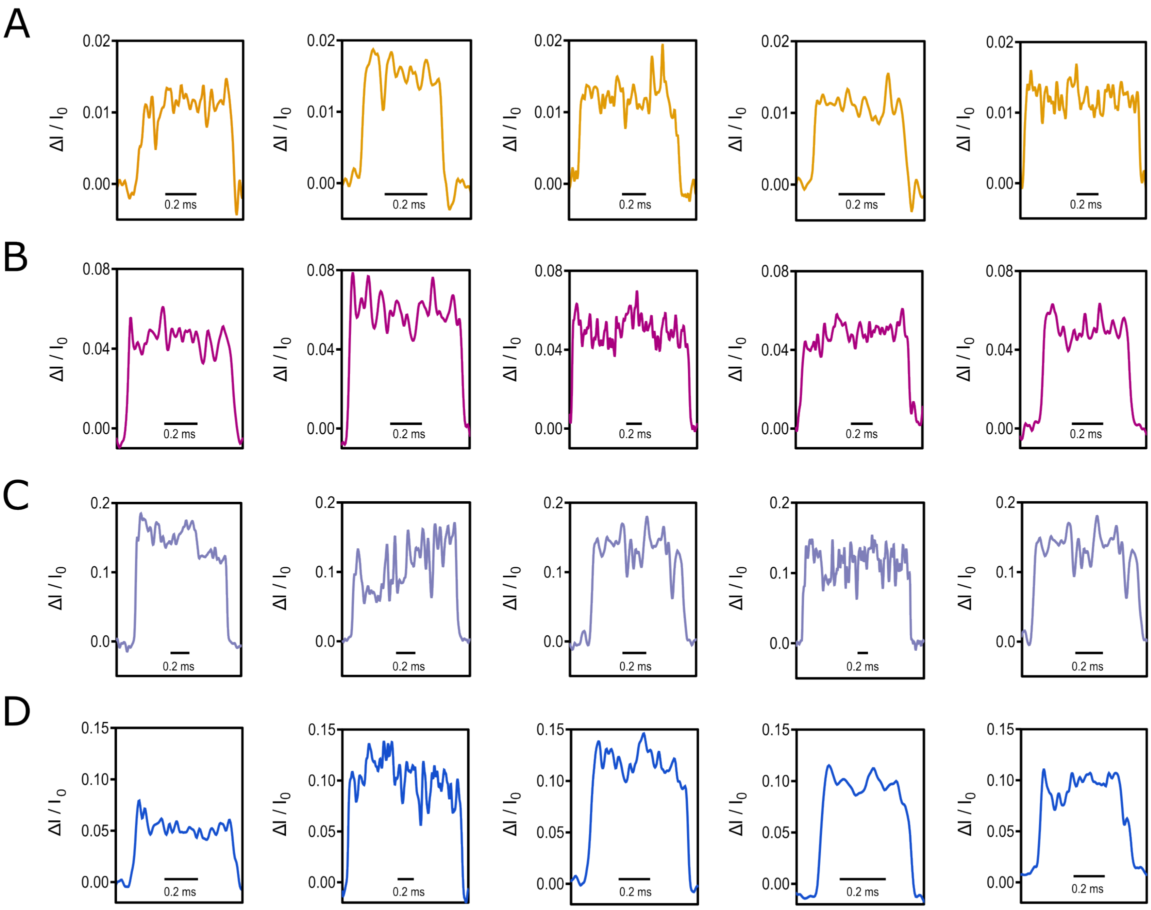

**Supplementary Figure S6. Representative individual resistive pulses** in the presence of (**A**) Fab, (**B**) IgG, (**C**) β-Galactosidase, (**D**) Apoferritin. The current recordings were collected using a recording buffer with 500 mM NaCl, 50mM Tris-HCl and 0.2 μM amphipol, pH 7.5 with a 200 kHz sampling rate and were filtered with a 10 kHz Gaussian low–pass filter for Fab and a 20 kHz Gaussian low–pass filter for IgG, β-Galactosidase and Apoferritin.

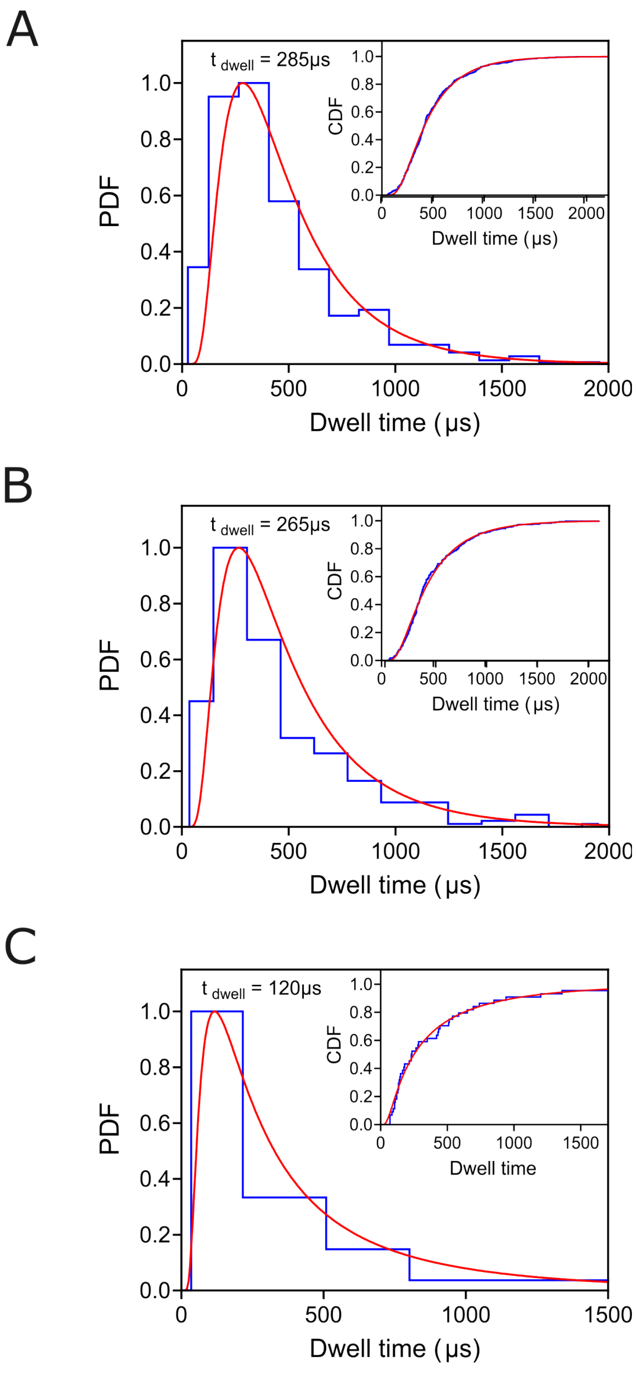

**Supplementary Figure S7. The distribution of dwell times $t_d$ of resistive pulses from** (**A**) Fab, with the most probable $t_d$ value of 285 μs. (**B**) IgG, with the most probable $t_d$ value of 265 μs. (**C**) β-Galactosidase, with the most probable $t_d$ value of 120 μs. The distributions include the $t_d$ values of all detected resistive pulses that were longer than 20 μs.

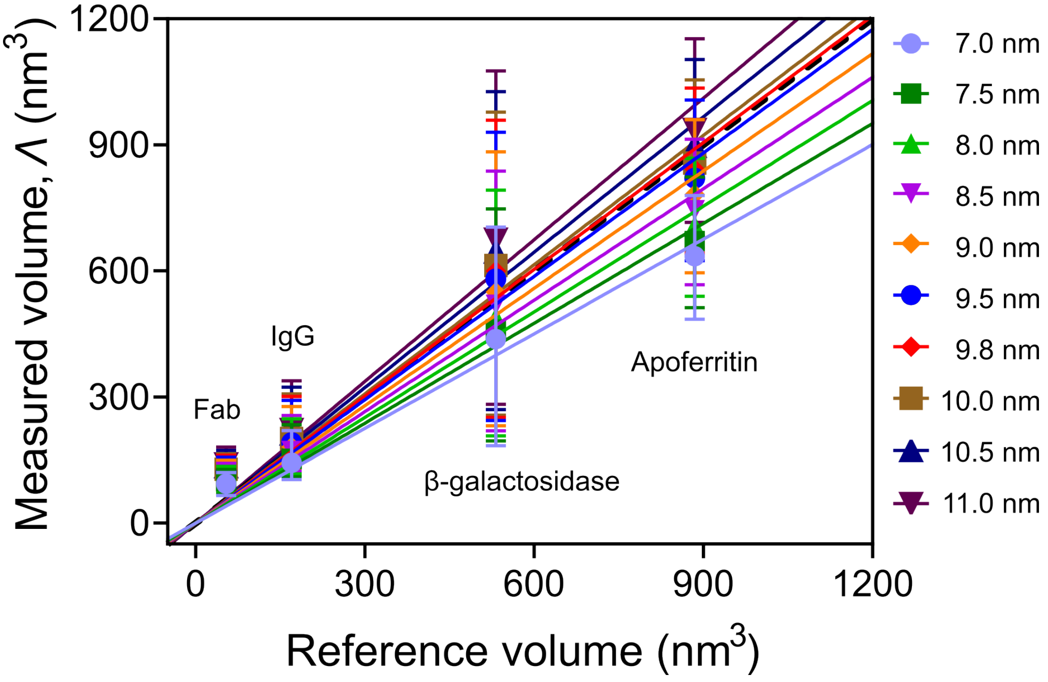

**Supplementary Figure S8. Empirical determination of pore length of PFO pores.** Comparison of excluded volumes measured in PFO nanopore experiments with the reference volume determined from the atomic coordinates (*.PDB files). We fitted linear regressions with a zero intercept to ensure accurate analysis. We first screened pore lengths from 7 to 11 nm in 0.5 nm increments to identify the range where possible pore length was likely to fall, then refined the search in 0.1 nm increments within that range (9.5 – 10 nm). The figure illustrates that a pore length of 9.8 nm most closely matches the ideal slope of 1.0 (black dotted line) of measured values as a function of the reference values of four different proteins.

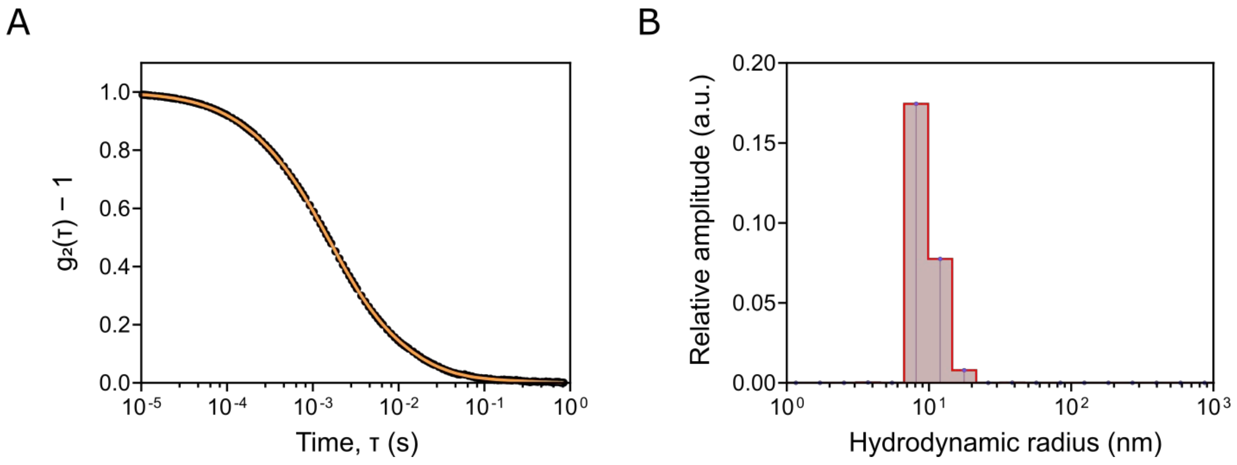

**Supplementary Figure S9. Dynamic light scattering (DLS) analysis of rAAV2 particles.** (**A**) Intensity autocorrelation function $g_2 - 1$ as a function of lag time obtained from DLS measurements of empty rAAV2, with the fitted curve shown in black. (**B**) Corresponding particle size distribution showing a monodisperse population with an average hydrodynamic radius of 10.9 ± 5.1 nm. The samples were prepared in a buffer containing 500 mM NaCl, 50 mM Tris-HCl, pH 7.5.

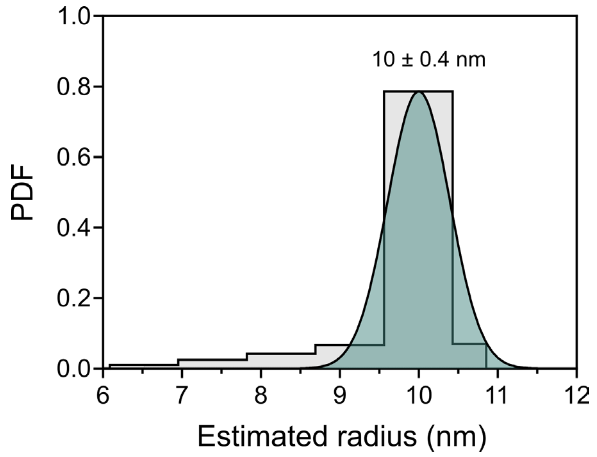

**Supplementary figure S10. Estimation of empty rAAV2 VLP radius.** Calculated radius of empty rAAV2 particles, obtained by converting the excluded volume from nanopore recordings to a radius using a spherical model, (see **Supplementary Note 4**), resulting in an estimated radius of 10 ± 0.4 nm.

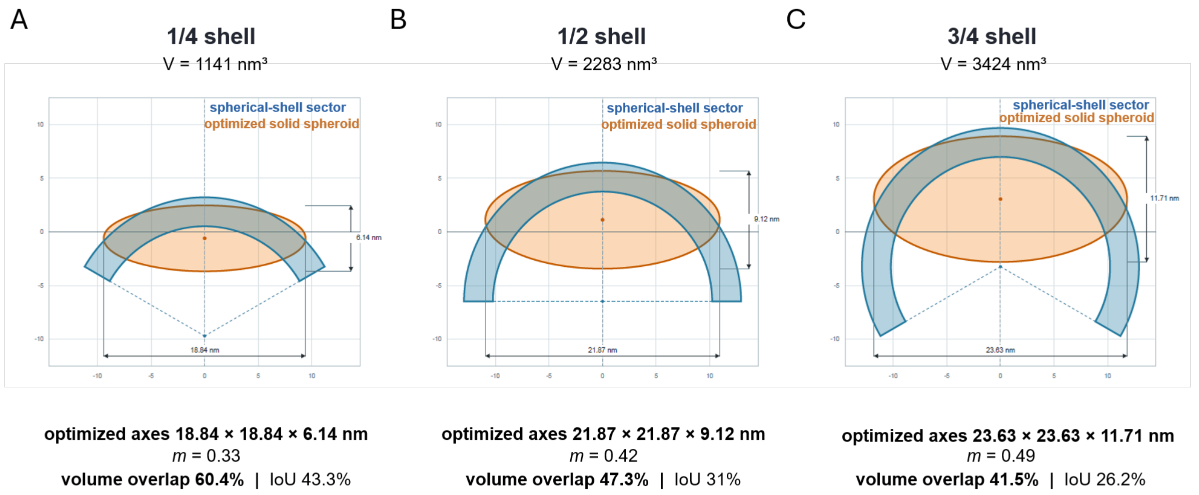

**Supplementary Figure S11. Modelling fragments of spherical shells of rAAV2 particles with optimized equal-volume ellipsoids of revolution.** To-scale meridional cross-sections compare rotationally symmetric spherical-shell fragments (blue) comprising 1/4, 1/2, and 3/4 of a complete shell with optimized solid oblate spheroids (orange). The complete shell had a volume of 4565 nm$^3$, an outer radius of 12.92 nm, an inner radius of 10.22 nm, and a uniform wall thickness of 2.70 nm, yielding fragment volumes of 1141, 2283, and 3424 nm$^3$, respectively. The shell fragments were defined as annular spherical sectors with polar half-angles of 60°, 90°, and 120°. For each fragment, the equatorial radius, polar radius, and axial position of an ellipsoid of revolution were varied while constraining its volume to equal that of the corresponding shell fragment. A global grid search followed by deterministic pattern refinement minimized the three-dimensional symmetric-difference volume between the shell and spheroid. Because the two objects had equal volumes, this procedure was equivalent to maximizing their intersection volume. Intersections were evaluated by exact analytical integration of their radial cross-sections; no constraint was imposed on the horizontal extent of the spheroid. IoU (intersection over union) quantifies the geometric similarity between the shell fragment and its fitted spheroid by comparing their shared volume with the total volume occupied by both shapes. The modelling workflow and computational script were developed with the assistance of ChatGPT using the “GPT-5.6 Sol Extra High” setting.

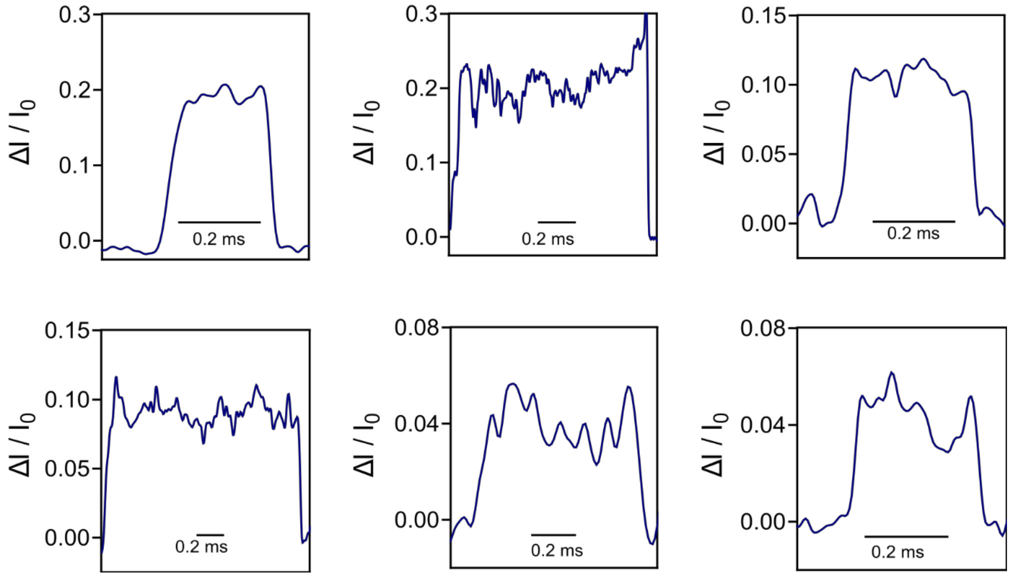

**Supplementary Figure S12. Representative individual resistive pulses in the presence of ribosome solution.** The current recordings were collected using 500 mM NaCl, 50 mM Tris-HCl, 10 mM $Mg(CH_3COO)_2$ and 0.2 µM amphipol at pH 7.5 with a 200 kHz sampling rate and were filtered with a 20 kHz Gaussian low-pass filter.

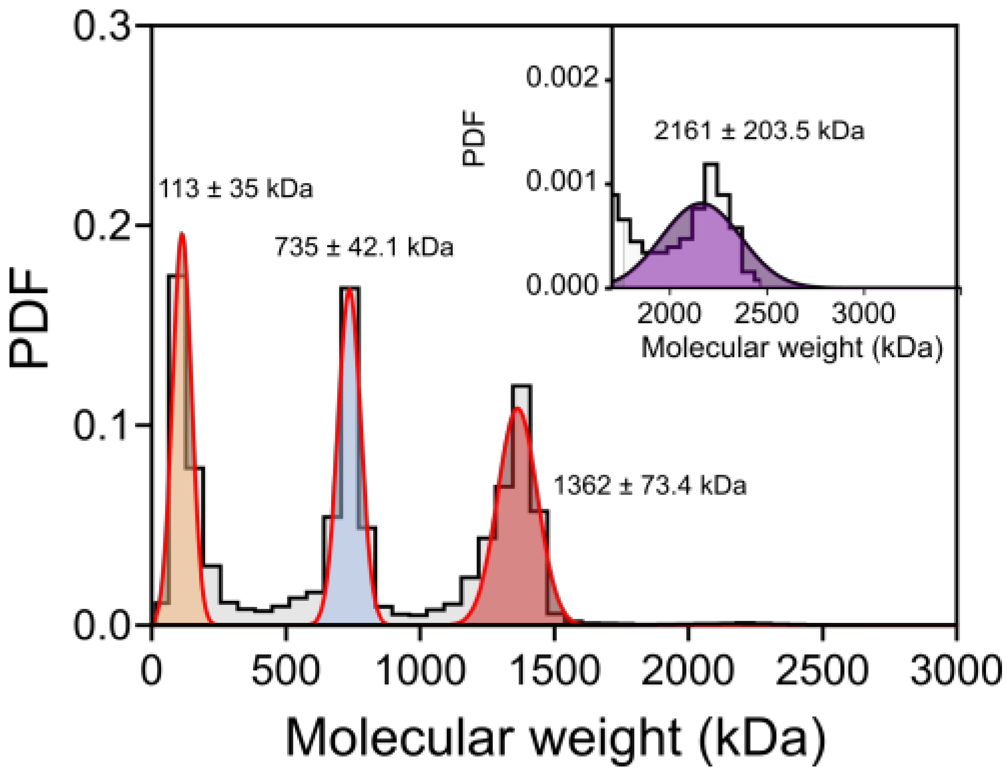

**Supplementary Figure S13.** Molecular weight distributions of ribosome solution determined by mass photometry in the recording buffer containing 500 mM NaCl, 50 mM Tris-HCl with a 0.2 µM amphipol pH of 7.5. A histogram was plotted using binning according to the Rice rule.

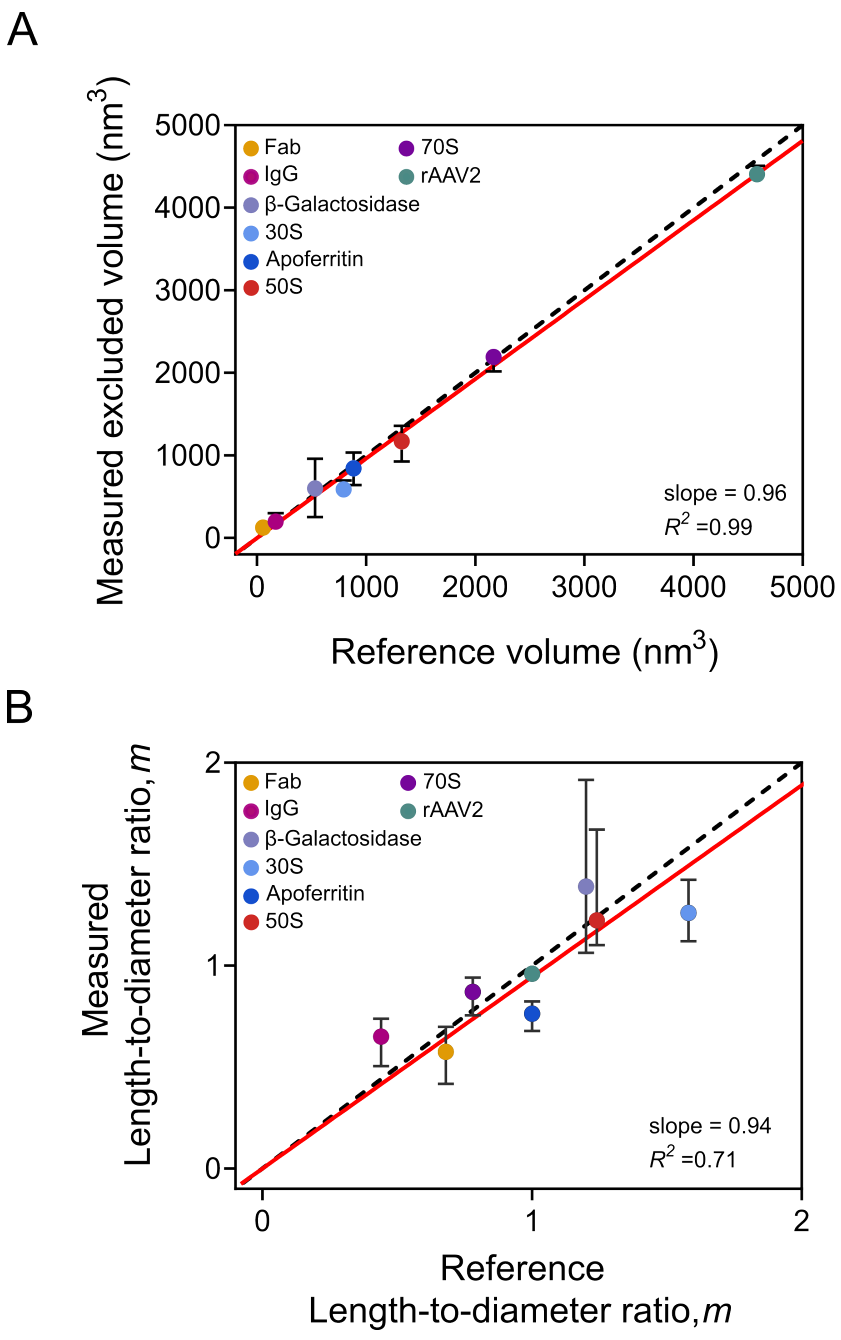

**Supplementary Figure S14. PFO nanopore-derived structural parameters across all analytes.** (A) Median values of the excluded volume (*Λ*) and (B) length-to-diameter ratio (*m*) determined from single-event analyses of all analytes are plotted against the theoretical reference values. Error bars show the first and third quartiles. The black dotted lines represent the ideal 1:1 agreement (slope = 1), and the red solid lines are the linear regressions performed imposing a zero intercept. For rAAV2, only particles with length-to-diameter ratio, *m,* between 0.9 and 0.99 are included in the analysis.

### *Supplementary Table*

**Supplementary Table S1. Estimation of excluded volume (*Λ*) and length-to-diameter ratio (*m)*** of all analytes used in this study, using molecular weight (MW)[10] of the protein, data analysis package[3] and PFO nanopore.

| Protein analyte | Referenced and measured excluded volume, Λ and length - to - diameter ratio, m | | |
|---|---|---|---|
| | Using MW of protein (Ref. 10) | Data analysis package (Ref. 9) | Measured via PFO |
| Fab | $\Lambda$ (nm$^3$) = 60.1 | $\Lambda$ (nm$^3$) = 54<br>m = 0.6 | $\Lambda$ (nm$^3$) = 125<br>m = 0.57 |
| IgG | $\Lambda$ (nm$^3$) = 180.1 | $\Lambda$ (nm$^3$) = 170<br>m = 0.44 | $\Lambda$ (nm$^3$) = 197<br>m = 0.65 |
| β-Galactosidase | $\Lambda$ (nm$^3$) = 558.6 | $\Lambda$ (nm$^3$) = 532<br>m = 1.2 | $\Lambda$ (nm$^3$) = 599<br>m = 1.38 |
| 30S | $\Lambda$ (nm$^3$) = 869 | $\Lambda$ (nm$^3$) = 794<br>m = 1.58 | $\Lambda$ (nm$^3$) = 586<br>m = 1.26 |
| Apoferritin | $\Lambda$ (nm$^3$) = 578.6 | $\Lambda$ (nm$^3$) = 885<br>m = 1 | $\Lambda$ (nm$^3$) = 841<br>m = 0.76 |
| 50S | $\Lambda$ (nm$^3$) = 1780 | $\Lambda$ (nm$^3$) = 1325<br>m = 1.24 | $\Lambda$ (nm$^3$) = 1246<br>m = 1.29 |
| 70S | $\Lambda$ (nm$^3$) = 2651 | $\Lambda$ (nm$^3$) = 2168<br>m = 0.78 | $\Lambda$ (nm$^3$) = 2270<br>m = 0.81 |
| rAAV2 VLP | $\Lambda$ (nm$^3$) = 4567 | *n.a* | $\Lambda$ (nm$^3$) = 4407<br>m = 0.96 |
